# WASp activity in macrophages prevents mechano-induced inflammation by protecting the nuclear envelope

**DOI:** 10.1101/2024.12.23.630061

**Authors:** Roberto Amadio, Giulia Bracchetti, Zahraa Alraies, Giulia Maria Piperno, Lucia Lopez Rodriguez, Mathieu Maurin, Martina Conti, Laura Andolfi, Maria Carmina Castiello, Francesca Ferrua, Anna Villa, Alessandro Aiuti, Ana-Maria Lennon-Dumenil, Federica Benvenuti

**Affiliations:** Cellular Immunology, International Centre for Genetic Engineering and Biotechnology (ICGEB), Trieste, Italy; University of Trieste, Department of Life Sciences (DSV), 34127 Trieste, Italy; INSERM-U932 Immunité et Cancer, Institut Curie, PLS Research University, 75005 Paris, France; Istituto Officina dei Materiali - National Research Council (CNR-IOM) - Basovizza, Trieste, Italy; San Raffaele Telethon Institute for Gene Therapy (SR-TIGET), IRCSS San Raffaele Scientific Institute, 20132 Milan, Italy; Institute for Biomedical Technologies - National Research Council (CNR-ITB), 20054 Segrate, Milan, Italy

**Author notes:** Laboratory of Cell and Molecular Biology, Experimental Research (C1), 005E - NYU Abu Dhabi, Saadiyat Campus, Abu Dhabi, United Arab Emirates. Corresponding author, Federica Benvenuti.

**Keywords:** Wiskott-Aldrich Syndrome, actin, macrophages, nuclear rupture, inflammation

## Abstract

Mutations in the immune-specific actin regulator WASp induce a proinflammatory state in myeloid cells, whose underlying causes remain poorly defined. Here, we applied microfabricated tools that mimic tissue mechanical forces to explore the role of WASp in connecting mechano-sensing to the activation of inflammatory responses in macrophages. We show that WASp-deficient macrophages carry alterations in nuclear structure and undergo increased blebbing and nuclear rupture when exposed to mechanical confinement. High-resolution imaging indicates that WASp drives the formation of protective perinuclear actin structures in response to confinement. Functionally, a proinflammatory gene signature linked to nuclear envelope rupture is preferentially active in confined WASp null macrophages, which partially depends on the cGAS-STING pathway of cytosolic DNA sensing. Analysis of transcriptional datasets of human and mouse tissue macrophages confirmed elevated inflammatory activation in WASp null cells. Together, these data uncover that WASp restricts pro-inflammatory activation of macrophages by preserving nuclear integrity in confined environments, providing novel clues to understand inflammatory activation in Wiskott-Aldrich Syndrome.

## Introduction

Tissue mechanical forces are emerging as key orchestrators of inflammation and immune responses in homeostatic and pathological conditions. Depending on the context, tissue rigidity and compression are transduced by diverse mechanisms to enhance or dampen cytokine production (Jain and Vogel, 2018; Du et al., 2022). Mechanical inputs are received by the nucleus, the larger and more rigid cellular organelle, through a complex connection between the nuclear lamina and the cytoskeleton, inducing morphological changes that regulate chromatin organisation and gene transcription (Nader et al., 2021b). However, errors in transmission or reception of mechanical cues may cause loss of nuclear envelope integrity and unwanted extrusion of chromosomal DNA into the cytosol (Härtlova et al., 2015; Dou et al., 2017; Dunphy et al., 2018), which is detected by cytosolic sensors such as cGAS and STING. Engagement of such receptors, triggers activation of cytokine genes, senescence or cell death, depending on the severity of damage (Dou et al., 2017; Mu et al., 2020; Nader et al., 2021a; Sladitschek-Martens et al., 2022), which may promote development of autoimmunity (Decout et al., 2021). Hence, maintaining nuclear envelope integrity is critical to restrict unwanted activation, particularly in immune cells that navigate complex environments, and are continuously exposed to mechanical compression (Denais et al., 2016; Raab et al., 2016; De Silva et al., 2023). Nuclear stability relies on a finely regulated balance between chromatin mechanics and cytoskeletal forces (Nava et al., 2020). Within cytoskeletal components, microtubules were shown to limit nuclear deformations in dendritic cells (Thiam et al., 2016), while intermediate filaments protect nuclei of murine embryonic fibroblast during constricted migration (Patteson et al., 2019). Arp2/3-dependent branched actin polymerization is critical to support nuclear deformation in migrating dendritic cells (Thiam et al., 2016), and it acts as a barrier to maintain nuclear integrity and limit DNA damage (Sladitschek-Martens et al., 2022; Delgado et al., 2024). Consistently, genetic ablation of the Cdc42 GTPase DOCK8 impairs nuclear integrity in T and dendritic cells during confined migration (Reis-rodrigues et al., 2024; Shen et al., 2024), suggesting that a broad range of diseases caused by mutations in regulators of actin branching may suffer from nuclear derangement.

The Wiskott-Aldrich Syndrome (WAS) is a primary immunodeficiency characterized by frequent autoimmune and autoinflammatory manifestations. WAS is caused by mutations in Wiskott-Aldrich Syndrome protein (WASp), a critical activator of the Arp2/3 complex which controls branched actin polymerization specifically in non-erythroid immune cells (Blundell et al., 2010; Naseem et al., 2022). WASp expression in immune cells regulates actin-driven processes such as cell migration (Gaertner et al., 2022; Oliveira et al., 2022), bacterial clearance (Lee et al., 2017), cytokine production (Prete et al., 2013; Cervantes-Luevano et al., 2018; Piperno et al., 2020; Amadio et al., 2021) and homeostasis of autoreactive B cells and Tregs (Sereni et al., 2018; Calixto Vieira et al., 2023). However, whether WASp plays a role in nuclear integrity is unknown. This may be particularly relevant in myeloid cells that experience continuous mechanical challenges in tissues and were shown to display and increased inflammatory phenotype in WAS.

In this work, we applied microfabricated tools that mimic tissues’ mechanical forces to address the role of WASp in preserving nuclear stability and inflammatory responses under mechanical pressure. By combining imaging and transcriptional profiling, we found that WASp-driven actin polymerization is necessary to counteract mechanical challenges and preserve nuclear envelope integrity in macrophages, restricting inflammatory activation.

## Results

### Nuclear defects in WKO macrophages

To explore the hypothesis of an altered nuclear phenotype in WAS, we started by analysing simple nuclear parameters in bone marrow derived macrophages derived from wild type or WASp null mice (WT and WKO hereafter) (Snapper et al., 1998; Prete et al., 2013). We assessed the area, roundness and presence of micronuclei in cells plated in 2D (Fig.1A). We found that the nuclei of WKO macrophages had an increased mean projected area, were more irregular and showed a higher incidence of micronuclei (Fig.1B), indicating general alterations in nuclear homeostasis. Previous findings in other systems linked defects in Lmna (encoding for Lamin A/C) and nuclear envelope folding to impaired mechanical tension and reduced protection from deformations (Kim et al., 2017; Tang et al., 2023). We thus focused on Lamin A/C and the nuclear envelope protein Emerin in our model. Total Lamin A/C levels were reduced in WKO, both at the protein and mRNA level, whereas mRNA expression of type B lamins and Emerin was not different (Fig.1C and 1D). Importantly, we noticed higher degree of Lamin A/C wrinkles and irregularities in nuclei of WASp null macrophages (Fig.1E), which coincided with Emerin accumulation (Fig.1F). Altogether these data suggest defective nuclear envelope homeostasis and increased folding of the nuclear envelope in the absence of WASp.

**Figure 1.**
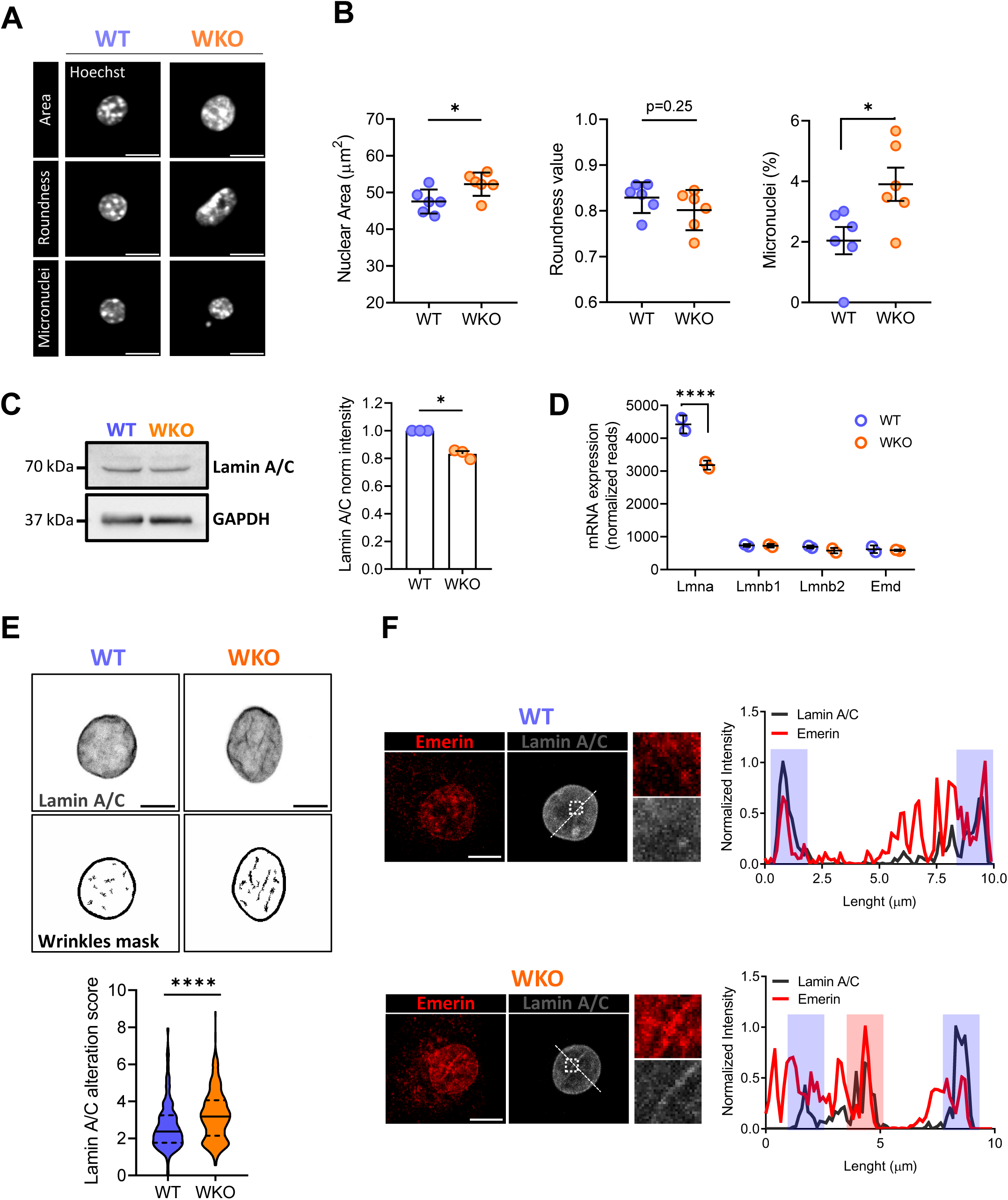
Nuclear parameters are modified in WASp null macrophages. (**A**) WT and WKO BMDM were plated on glass and labelled with Hoechst to visualise nuclei. Representative images show examples of the three parameters analysed: area (top), roundness (middle) and micronuclei (bottom). Scale bar 10 µm. (**B**) Quantification of the three nuclear parameters depicted in A, each dot is an independent experiment (*N*=6 experiments with *n*>2000 nuclei analysed for each genotype). Two-tailed unpaired t-test; \**P*<=0.05. (**C**) Representatives immunoblot (top left) and relative quantification of LMNA/C protein expression relative to GAPDH (top right); *n*=3 biological replicates. Data were analysed by one sample t-test. \**P*<=0.05. (**D**) Expression of lamin isoforms from RNA-seq analysis of WT and WKO BMDM. Normalized reads for Lmna, Lmnb isoforms 1 and 2 and Emerin. Representative data from *n*=2 replicates. Data were analysed by two-way ANOVA test with multiple comparisons and Sidak’s correction method. \*\*\*\**P*<=0.0001. (**E**) Representatives maximum z-projection of Lamin A/C staining (top) and the corresponding segmentation mask for wrinkles detection (bottom). Quantification of Lamin A/C alterations as defined in methods. A higher score indicates less homogeneity and more wrinkles. *n*=402 WT and *n*=302 WKO individual nuclei measured, in 3 independent experiments. Scale bar 5 µm. Two-tailed unpaired t-test, **** = *P*<0.0001. (**F**) Representative images of Lamin A/C (grey) and Emerin (red) co-staining (top) and line intensity profiles (bottom). Histograms depict areas of colocalization at nuclear boundaries (blue area) and in nuclear wrinkles (red area). Three independent co-labelling experiments were performed. Scale bar 5 µm.

Next, as the nucleus is highly connected to chromatin and cellular mechanics, we explored chromatin parameters and global cellular stiffness. We did not observe substantial differences in global heterochromatin content and peripheral heterochromatin between the two genotypes by chromocenters analysis (Supplemental Fig.1A). Similarly, no changes in heterochromatin distribution were detected by H3K9me2 staining, but we observed an increase in the total intensity of this marker (Supplemental Fig.1B). Cellular mechanics, as measured by Atomic Force Microscopy (AFM) indentation (Supplementary Fig.1C) showed that WKO cells were stiffer than WT ones (Supplementary Fig.1D). Of note, a stiffer cytoplasm has been linked to reduced nuclear stability under 3D physical constrains (Bastianello et al., 2023).

### Increased constricted migration and nuclear deformation in WKO macrophages

Next, we addressed the role of WASp in controlling the mechanical properties of the nucleus. First, we performed an assay to force nuclear deformations inside narrow constrictions using microchannels (Vargas et al., 2016). We adopted different channel dimensions, ranging from 2 µm to 8 µm in width and having a constant height of 5 µm. Macrophages were seeded in the pre-channel chamber and recorded overnight. The number of cells that entered the channel was calculated at the end of the recording period by counting the number of cells inside channels and normalizing by the number of approaching cells (Supplementary Fig.2A; materials and methods). As expected, reducing the size of the channels inhibited the entrance of cells inside constrictions (Fig.2A). Interestingly, the entrance rate of WKO macrophages was consistently higher, especially at smaller constrictions (Fig.2B). By carefully inspecting nuclear shape in cells inside channels we observed enhanced frequencies of nuclear herniations in WKO (Fig.2C). Herniations were preferentially observed at the front of the cell in the direction of migration (Fig.2D), in line with previous observations in other cell types (Denais et al., 2016). Since herniations predispose to nuclear rupture, we next transduced macrophages with a lentiviral construct expressing a nuclear localization signal (NLS)-tagged GFP (NLS–GFP), which is retained inside the nucleus and it is released upon nuclear envelope (NE) rupture (Fig.2E). Interestingly, we observed a larger fraction of NE rupture and cytosolic release of fluorescent signal in WKO macrophages than in WT (Fig.2F and 2G). Moreover, and consistently with increased nuclear fragility, we detected multiple rupture events on the same cell exclusively in WKO (Fig.2H). A parallel set of transwell migration experiments confirmed the superior ability of WASp null cells to translocate through small pores (Supplementary Fig.2B). We ruled out that enhanced transwell migration in WKO may depend on changes in chemokine receptors expression (Supplementary Fig.2C). Notably, migration induced an increase in nuclear deformation and enhanced γH2AX staining, particularly in WKO cells (Supplementary Fig.2D). Overall, these data indicate that WASp-deficient macrophages are facilitated in entering small constriction, in line with a previous report on dendritic cells (Gaertner et al., 2022), suggesting reduced resistance to nuclear deformation. Importantly, the nuclear envelope of WKO cells ruptures during constricted migration, releasing nuclear material into the cytosol.

**Figure 2.**
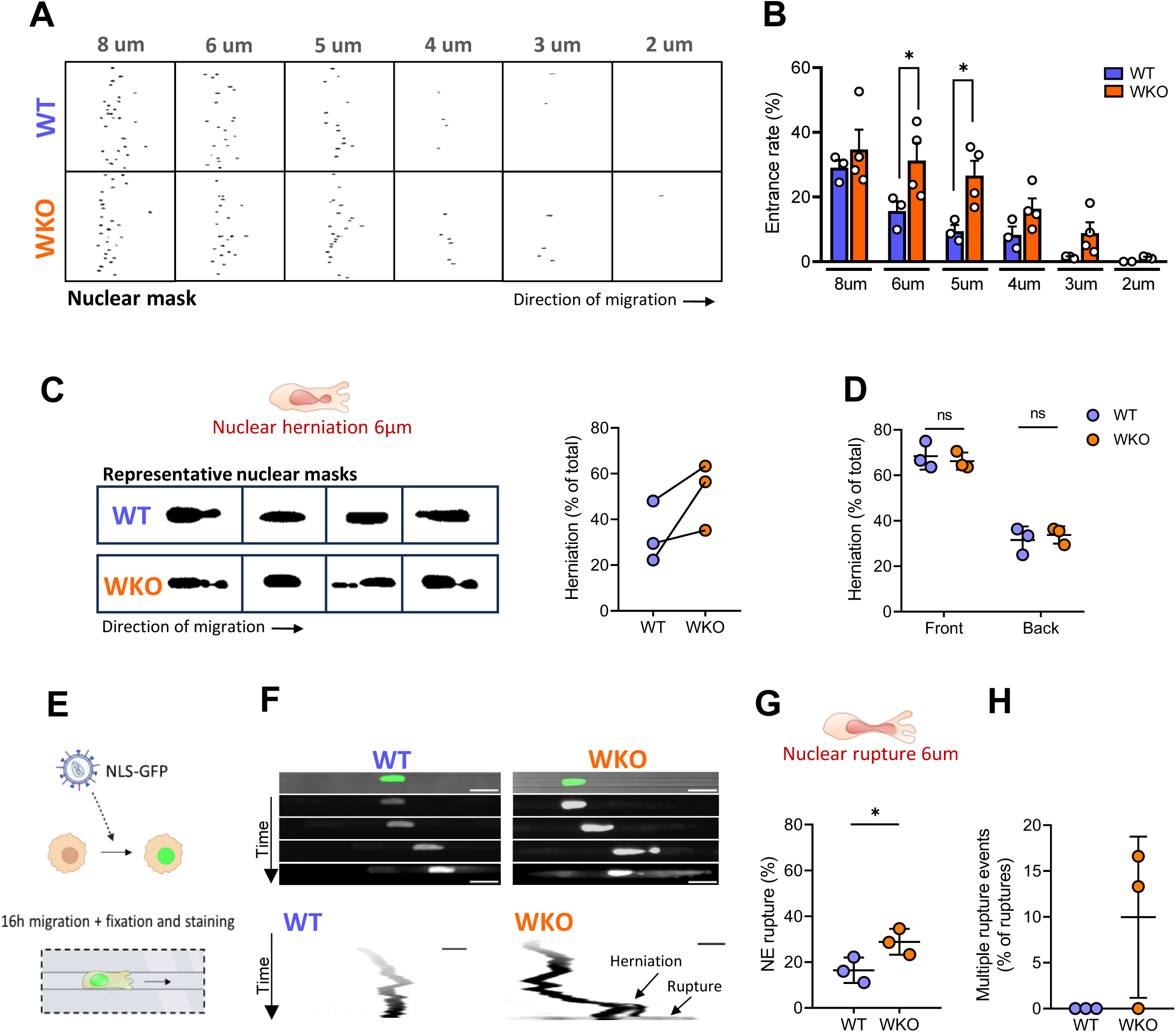
WASp controls the deformation and integrity of the nucleus during migration in microchannels. (**A**) Analysis of nuclear parameters in macrophages migrating through 2-8 μm x 5 μm fibronectin coated micro-channels. Representative images of nuclei within channels after overnight migration (applying a nuclear mask). (**B**) Entrance rate of the two genotypes inside channels of different dimensions. Data are from *N*=3 and *N*=4 independent experiments for WT and WKO, respectively. Data were analysed by two-way ANOVA test, \**P*<=0.05. (**C**) Representative examples of WT and WKO macrophage nuclei inside 6 µm x 5 μm channels (left). Corresponding quantification of the fraction of cells showing herniated nuclei (right). *n*=40 WT and 57 WKO cells in *N*=3 independent experiment, two-tailed unpaired t-test. (**D**) Quantification of the position of the herniation relative to the direction of migration, two-way ANOVA with Sidak’s correction method. ns, not significant. (**E**) WT and WKO BMDM were transduced with NLS-GFP to track nuclear rupture during migration. (**F**) Representative time frames (top) and relative kymographs (bottom) showing a nuclear rupture event in WKO migrating inside 6 µm x 5 μm channels, detected as cytosolic leakage of NLS-GFP. Scale bar 10 µm. (**G**) Quantification of the fraction of cells undergoing nuclear rupture during live imaging. *N*=3 independent experiments, with at least 20 individual cells analysed per genotype for each experiment. Two-tails unpaired t-test. \**P*<=0.05. (**H**) Quantification of multiple nuclear ruptures detected during live imaging experiments expressed as fraction of total ruptures recorded.

### WASp limits nuclear ruptures under mechanical confinement

To more faithfully recapitulate deformation events experienced by macrophages in tissues, we next applied a vertical confiner device (Alraies et al., 2024) based on PDMS micropillars (Supplementary Fig.3A). The heights of BMDM nuclei plated on fibronectin-coated glass slides without confinement range between 5 and 6 µm (Supplementary Fig.3B), therefore we selected 6 and 3 µm height as mild and strong confinement, respectively (Supplementary Fig.3C). Confocal images were acquired after one hour of confinement to examine nuclear parameters. We classified nuclei into two categories: 1) intact, when the nuclear boundaries were regular and continuous; 2) damaged, when blebs or herniations were found to interrupt the nuclear perimeter (Fig.3A). The fraction of damaged nuclei in WASp null cells was slightly higher under mild confinement and significantly higher under strong confinement (Fig.3B), suggesting loss of nuclear integrity. Indeed, measuring NLS-GFP localization after confinement at 3 µm, we observed an increased frequency of NE rupture and NLS-GFP cytosolic release in WKO cells (Fig.3C and 3D). Live tracking of NLS-GFP leakage revealed that rupture correlates to blebbing/herniation of the nuclear surface (Fig.3E). To causally link WASp to nuclear deformation and loss of NE integrity, we next transduced WASp null macrophages with a lentiviral construct expressing WASp-GFP (Supplementary Fig.3D) (Blundell et al., 2008). This resulted in around 50% of cells expressing a clear GFP signal and WASp-GFP fusion protein of the expected size (Supplementary Fig. 3E and 3F). The resulting mixed population of GFP-positive (rescued) and negative (control) cells were subjected to confinement at 3 µm. In this highly controlled setting, we found that GFP-positive cells showed significantly fewer nuclear blebs than GFP-negative ones (Fig. 3F and 3G). Moreover, the fraction of cells showing Lamin A/C rupture was reduced in WASp-GFP positive cells (Fig.3H and 3I), confirming a direct impact of WASp in preserving nuclear stability under confinement.

**Figure 3.**
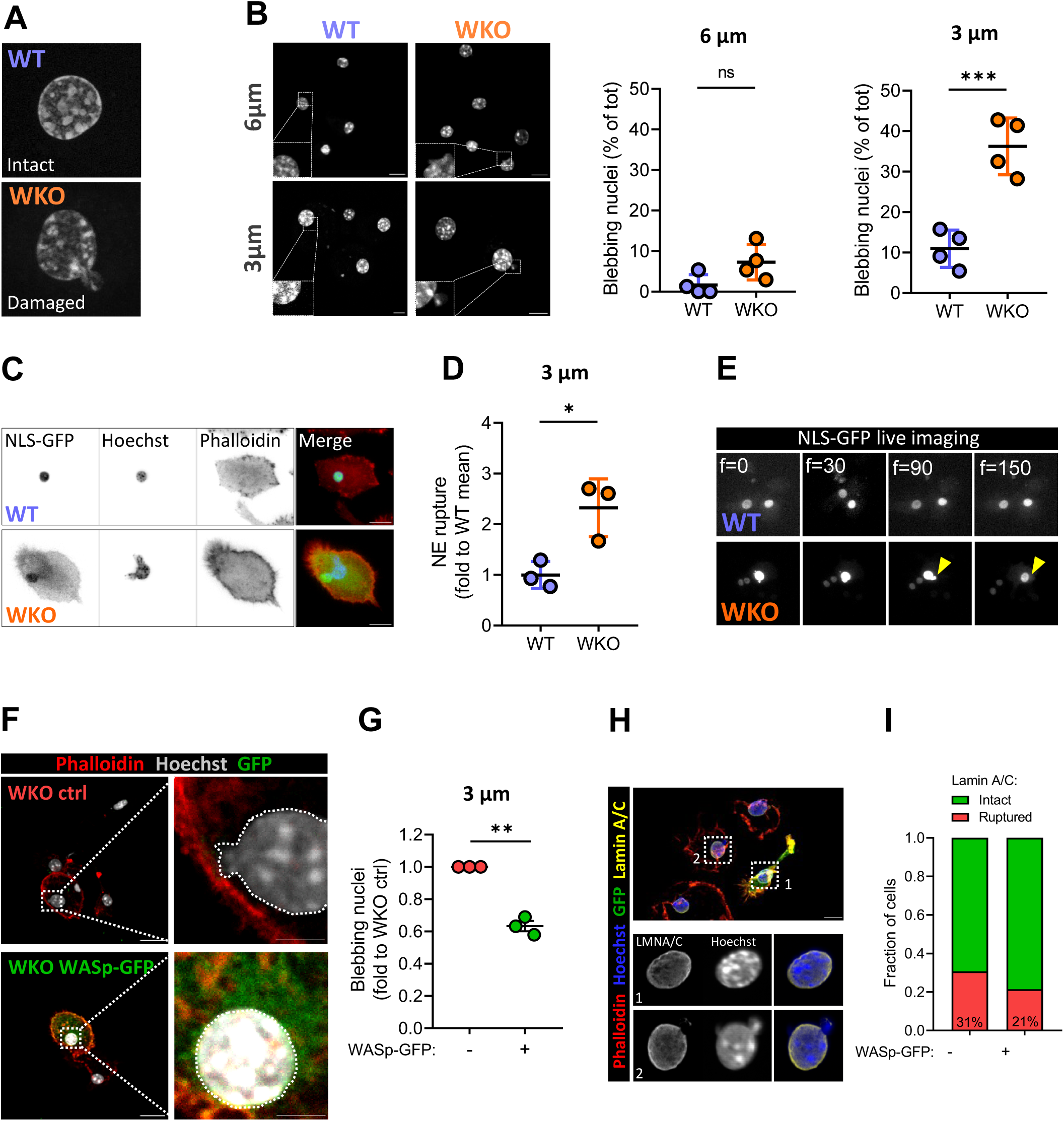
Vertical confiners induce higher deformation and rupture in WASp-deficient macrophages. (**A**). Examples of intact or damaged nuclei, as used for quantification. (**B**) Representative images of WT and WKO macrophage’s nuclei under 3 or 6 μm confinement height (right). Insets show nuclear deformation. Scale bar 10 µm. Quantification of the fraction of cells carrying a damaged nucleus after 1 h confinement at the indicated height (left). At least 200 cells per conditions were analysed in *N*=3 independent experiments. Two-tailed unpaired t-test. \*\*\**P*<=0.001. (**C**) Assay on NLS-GFP cytosolic release under confinement. Representative images of NLS-GFP^+^ macrophages after 1h of confinement at 3 µm. Scale bar 20 µm. (**D**) Corresponding quantification of nuclear ruptures upon confinement based on NLS-GFP cytosolic leakage in C. Each dot is an independent experiment (*N*=3). *n*=173 WT and *n*=102 WKO cells. Data are expressed as fold to WT mean. \**P*<=0.05 in a two-tailed unpaired t-test. (**E**) Individual frames from a live imaging experiment showing nuclear envelope rupture during following bleb formation in WKO (yellow arrow). The timeframe is indicated by f. (**F**) Representative images of WASp-GFP^+^ and WASp-GFP^-^ macrophages after 1 h confinement at 3 µm. Dashed lines mark nuclear boundaries. Scale bar 20 µm; 5 µm for insets. (**G**) Corresponding quantification of nuclear blebbing in control or WASp-GFP reconstituted cells. Data are expressed as fold on WKO ctrl, *n*=83 WT and *n*=99 WKO cells from *N*=3 independent experiments, one sample t-test. \*\**P*<=0.01. (**H**). Representative images of confined (3μm) WASp-GFP^+^ and WASp-GFP^-^ macrophages stained for Lamin A/C (yellow). Nuclear insets show Lamin A/C and Hoechst signal in WKO ctrl (1) and WKO WASp-GFP (2). Scale bar 20 µm. (**I**) Quantification of Lamin A/C rupture events in WKO ctrl and WKO WASp-GFP based on **H**. Data are from n=135 WKO ctrl and n=22 WKO WASp-GFP in *N*=2 experiments.

### WASp directs perinuclear actin polymerization in confined macrophages

We then aimed to investigate a potential mechanism by which WASp and actin polymerization support nuclear stability during mechanical challenges. We took advantage of a fixation-under-confinement protocol, which preserves cytoskeletal structures once confinement is released (Alraies et al., 2024). We found that confinement triggered the accumulation of dense actin structures in the central body of the cell in WT, but not in WKO macrophages (Fig.4A). These actin patches closely resemble the Arp2/3-dependent structures described in mechanically confined dendritic cells with a role in nuclear envelope unfolding (Alraies et al., 2024). By high-resolution imaging, we further classified confined cells into four categories according to actin distribution: 1) perinuclear actin patches; 2) intranuclear actin accumulation; 3) filamentous perinuclear actin ring and 4) diffuse cytosolic actin (Fig.4B). Of note, perinuclear actin patches were observed in a substantial proportion of WT cells but were virtually absent in WKO (Fig.4C and 4D). Intranuclear actin accumulation was present in a small fraction of cells, in similar frequencies between the two genotypes. Additionally, a small fraction of WKO cells showed thin filamentous actin surrounding the nucleus (Fig.4C and 4D), a phenomenon not detected in WT macrophages. To explore the role of WASp in driving the polymerization of perinuclear actin structures, we leveraged WKO macrophages transduced with WASp-GFP for colocalization analysis (Fig.4E). Interestingly, the Mander’s coefficient between phalloidin and GFP was high in the perinuclear actin patches induced by confinement (Fig.4F), strongly suggesting an active role of WASp in directing the formation of these structures. Consistently, also the distribution of WASp-GFP in macrophages inside microchannels was enriched in the central part of the cell body and around the nucleus (Supplementary Fig.4A and 4B). We conclude that WASp drives the nucleation of perinuclear actin patches in macrophages exposed to mechanical confinement. Together with the previous observations, this result suggests that a local perinuclear actin response may help to preserve nuclear integrity.

**Figure 4.**
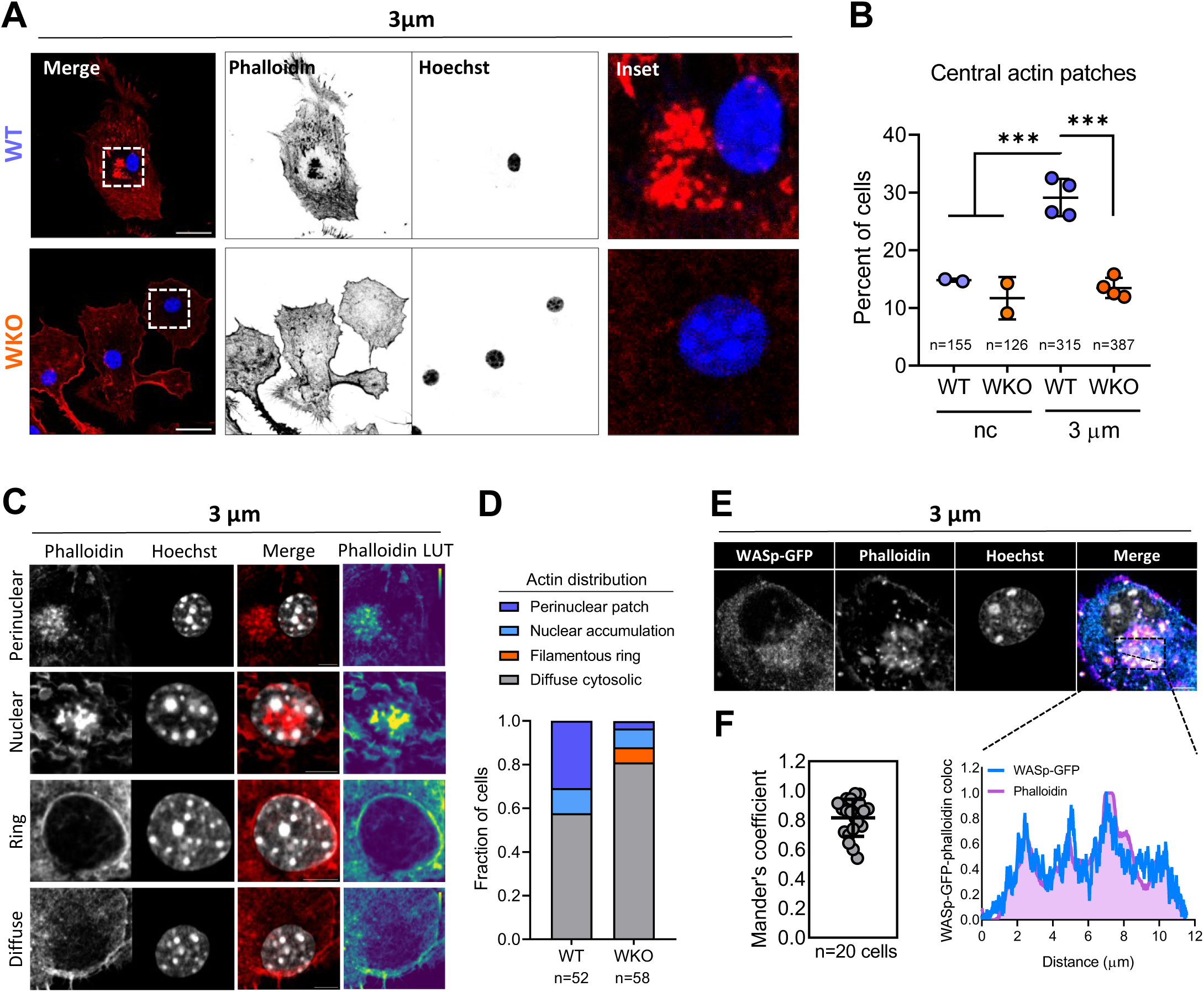
WASp triggers perinuclear actin polymerization under confinement. (**A**) Representative images of actin distribution in WT and WKO BMDM confined at 3 µm height. Scale bars 20 µm. (**B**) Quantification of the fraction of cells with central actin accumulation. n=number of cells, as indicated in figure. *N*=2 experiments for unconfined controls and *N*=4 for confined samples. Two-tailed unpaired t-test. \*\**P*<=0.01. (**C**) Representative images to define patterns of actin distribution in macrophages confined at 3 μm. Scale bar 5 µm. (**D**) Corresponding quantification of the fraction of WT and WKO macrophages in each category. n=number of cells, as indicated in figure; from *N*=3 individual experiments. (**E**) WKO cells transduced with WASp-GFP were confined at 3 µm and analysed to visualize the relative distribution of WASp and F-actin. Scale bar 5 µm. Line intensity profile of selected region showing the intensity distribution of WASp-GFP and phalloidin (bottom). (**F**) Quantification of WASp-GFP and F-actin Mander’s coefficient of colocalization on perinuclear actin patches. Data are from **E**, with a total of *n*=20 cells analysed from *N*=2 independent experiments.

### Confinement-induced nuclear ruptures exacerbate inflammation in WKO macrophages

To investigate the functional consequences of increased nuclear fragility in WASp null macrophages, we then analysed transcriptional changes induced by confinement (Fig.5A). WT and WKO macrophages at steady state (unconfined) express a common transcriptional program and some genotype-specific genes (Supplementary Fig.5A). Remarkably, WKO cells were enriched in immunoglobulin coding genes, an observation that deserves future investigations. Moreover, transcripts related to extracellular matrix organization, vascular remodelling and cellular activation/migration were higher in unconfined WKO macrophages than in WT. In contrast, genes controlling cellular metabolism were less expressed in WKO (Supplementary Fig.5A and Supplementary Table 1). Confinement induced profound transcriptional remodelling in both genotypes (Supplementary Fig.5B). Analysis of differentially expressed genes (DEGs) between unconfined and confined cells unveiled a core of 130 genes commonly upregulated by confinement in WT and WKO (Fig.5B), including some inflammatory cytokines and chemokines (Fig.5C). In addition, we observed 151 genes upregulated only in confined WKO cells and 36 in WT cells. Interestingly, genes selectively upregulated in WKO were related to inflammatory activation and tissue remodelling (Fig.5B), paralleled by reduced expression of metabolic pathways (Supplementary Fig.5C).

**Figure 5.**
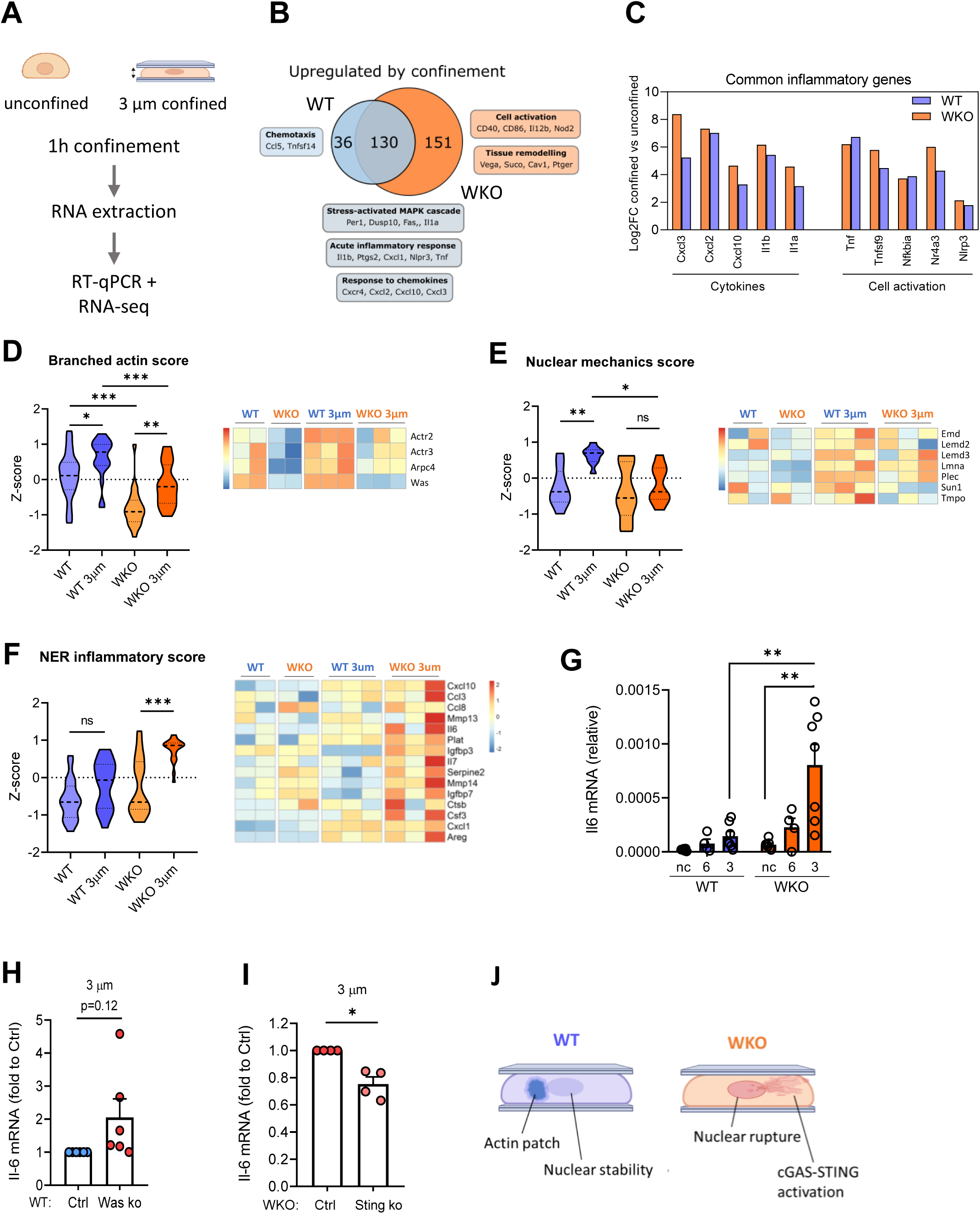
Mechanical confinement triggers innate activation in WASp KO macrophages. (**A**) BMDM were subjected to 1 h confinement at 3 µm or kept in unconfined conditions and processed for RNA-seq and gene qPCR expression analysis. (**B**) Venn diagram of common and unique upregulated DEGs in WT and WKO BMDM confined at 3 µm as compared to their respective unconfined controls. Relevant processes and genes are highlighted. (**C**) Log2FC of commonly upregulated inflammatory genes by confinement in the two genotypes. (**D**) Z-score of branched actin signature across the two genotypes in unconfined and confined conditions (left). Normalized expression of relevant individual genes is showed as heatmap (right). (**E**) Z-score of nuclear mechanic signature across the two genotypes in unconfined and confined conditions (left). Normalized expression of relevant individual genes is showed as heatmap (right). (**F**) Z-score of NER induced inflammatory signature across the two genotypes in unconfined and confined conditions (left). Normalized expression of relevant individual genes is showed as heatmap (right). *P* in **D**, **E**, and **F** were calculated by one-way ANOVA test. \**P*<=0.05, \*\**P*<=0.01, \*\**P*<=0.001. (**G**) Quantification of Il6 mRNA in confined WT and WKO BMDM after 1 h at indicated heights. *n*=6 for unconfined and 3 µm confined samples, *n*=4 for 6 µm confined samples. \*\**P*<=0.01 on a one-way ANOVA test. (**H**) Normalized expression of Il6 mRNA in WT ctrl and WT WASp KO BMDM. Data from 6 individual confinement replicates. (**I**) Normalized expression of Il6 mRNA in WKO ctrl and WKO STING KO BMDM from 4 individual confinement experiments. \**P*<=0.05 on a one-sample t-test. (**J**) Model of WT and WKO macrophage response to confinement.

To explore the molecular mechanisms underlying cellular responses to confinement, we evaluated the expression of genes implicated in cytoskeletal dynamics and control of nuclear stability (Supplementary Table 2), across all the conditions. A signature encompassing genes that control actin branching (branched actin score) was significantly higher in WT than in WKO macrophages, suggesting that WASp deficiency broadly impacts this process. Upon confinement, this gene set was upregulated in both genotypes, indicating fast cellular adaptation to mechanical inputs (Fig.5D), however it remained lower in WKO cells. This trend was even more prominent when selecting specific WASp effectors that have been directly linked to NE integrity (*Actr2*, *Actr3*, *Arpc4*) (Sladitschek-Martens et al., 2022; Delgado et al., 2024); showing a drastic reduction of expression in both unconfined and confined WASp deficient cells. Of note, a second gene module linked to nuclear mechanics including several genes previously implicated in NE stability, such as Lamin A/C, Emerin and Lap2β (Tmpo) (Chen et al., 2021; Lavenus et al., 2022; De Silva et al., 2023) was significantly upregulated in WT cells, but not in WKO, upon confinement (Fig.5E). Conversely, SUN2 and RhoA/ROCK, whose overexpression has been linked to nuclear blebbing and innate activation (Mu et al., 2020), were higher in WKO cells, together with RAC2-related genes (Supplementary Fig.5D). All together, these data suggest a defective branched actin-based response to confinement in WKO macrophages, concomitant to induction of a compensatory RhoA/RAC-mediated pathway, with a potentially negative impact on nuclear stability.

Defective mechanotransduction and reduction in the expression of genes linked to actin-dependent nuclear integrity were previously shown to induce activation of a proinflammatory cell state in Arp2/3 deficient fibroblast (Sladitschek-Martens et al., 2022). Hence, we examined the expression of this signature, defined as “nuclear envelope rupture (NER) inflammatory score” in our dataset. Remarkably, the NER inflammatory score was selectively activated in confined WASp null macrophages, causally connecting WASp to the control of proinflammatory activation in cells exposed to mechanical inputs (Fig.5F). Among the genes included in NER inflammatory score, Il6 was selected to perform further functional analyses. qPCR confirmed that Il6 transcripts were preferentially induced by confinement in WKO (Fig.5G). To exclude any potential bias in macrophages derived from constitutive WASp KO animals, we induced acute WASp deletion in WT BMDM by CRISPR-Cas9 gene editing (Supplementary Fig.5E). Importantly, WASp-depleted macrophages responded to confinement by augmented Il6 induction as compared to control cells (Fig.5H), indicating that WASp directly restricts confinement-induced activation. Given the widespread implications of the cGAS-STING pathway of cytosolic DNA sensing (Ablasser and Gulen, 2016) to innate activation and specifically in driving the NER inflammatory score (Sladitschek-Martens et al., 2022), we assessed its contribution to confinement-induced activation of WKO cells. STING ablation in WKO macrophages (Supplementary Fig.5F) blunted Il6 production upon direct stimulation by DMXAA, confirming the functional disruption of the pathway (Supplementary Fig.5G). We then exposed WKO macrophages carrying or not STING depletion to confinement at 3 µm of height.

Induction of Il6 transcripts was significantly reduced in WKO STING KO macrophages as compared to STING-proficient cells, causally connecting the pathway to the proinflammatory state of WKO macrophages (Fig.5I). Nevertheless, the uncomplete inhibition of Il6 expression in STING-depleted cells suggests that other sensors/mechanisms may contribute to the inflammatory activation downstream of NE ruptures in WASp-deficient cells. We conclude that increased NE rupture in WASp null macrophages triggers the expression of a specific proinflammatory signature (Fig.5J).

### Inflammatory phenotype in mouse and human WASp deficient macrophages

To validate our findings *in-vivo* we next focused on lung alveolar macrophages (AMO), a representative population of long-lasting resident macrophages constantly exposed to physical constraints and subjected to almost 2D confinements (De Silva et al., 2023). Flow cytometry profiling of AMO (Supplementary Fig.6A), showed higher levels of the activation marker CD86 in WKO mice (10 weeks of age), as compared to WT (Fig.6A). We next isolated AMO from 10 weeks and 25 weeks old mice. The absolute number of AMO was higher in WKO as compared to controls, especially at 25 weeks of age (Fig.6B). Notably, Il6 transcripts were slightly elevated in young animals and significantly higher in AMO from 25 weeks old WKO mice (Fig.6C). Therefore, we conclude that WASp is dispensable for differentiation or survival of lung tissue resident macrophages, however it controls their homeostatic inflammatory state.

**Figure 6.**
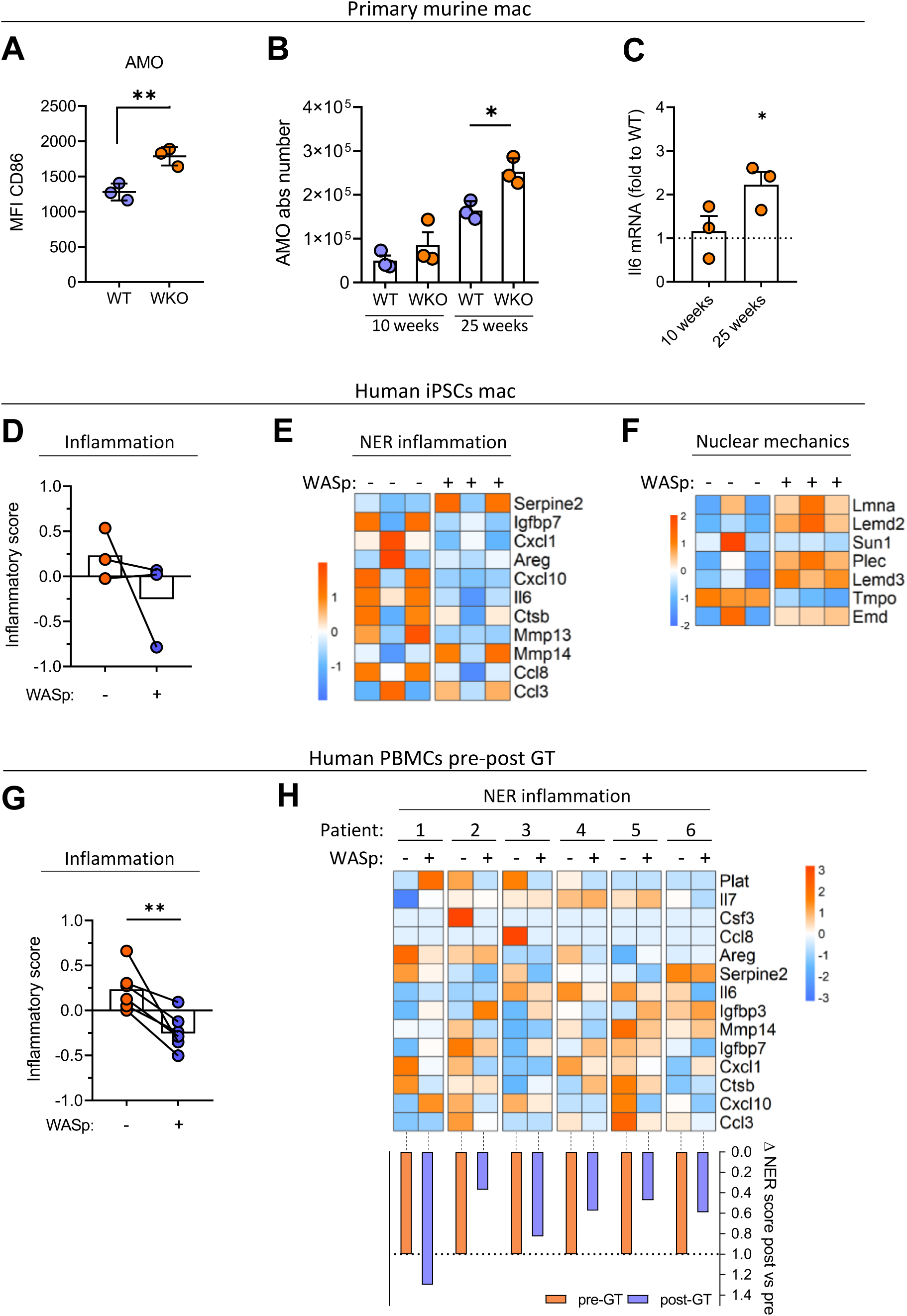
WASp deficiency correlates to enhanced expression of inflammatory genes in macrophages. (**A**) Alveolar macrophages (AMO) from 10 weeks old mice were harvested and processed for flow cytometry. Data show mean fluorescent intensity (MFI) of CD86 from one representative of two experiments (*n*=3 mice per genotype). Two-tailed unpaired t-test; \*\**P*<=0.01. (**B**) Quantification of AMO total numbers in 10- and 25-weeks-old adult mice (*n*=3 mice, data are from one representative of two experiments). One-way ANOVA test with multiple comparisons, \**P*<=0.05. (**C**) Il6 expression in AMO. mRNA expression (RT-qPCR, plotted as fold to WT) of the indicated genes in sorted AMO from 10-and 25-weeks-old WKO mice. *n*=3 mice for each condition, one sample t-test, \**P*<=0.05. (**D**) Expression of inflammatory signature in WASp proficient (WASp +) or WASp null (WASp -) human iPSCs-derived macrophages from GSE107963 dataset. (**E**) Heatmap of NER inflammatory genes expression (Log2+1 TPM normalized expression, z-score gene-wise scaled) of WT and WASp null iPSCs-derived macrophages from GSE107963. (**F**) Heatmap of genes related to nuclear mechanics (Log2+1 TPM normalized expression, z-score gene-wise scaled) of WT and WASp null iPSCs-derived macrophages from GSE107963. (**G**) Inflammatory score in PBMCs of a cohort of 6 WAS patients before (WASp -) and after restoration of WASp by gene therapy (WASp +, post-GT). (**H**) Heatmap of NER inflammatory genes expression (Log2+1 normalized reads, z-score gene-wise scaled) in human PBMCs from WAS patients pre- and post-gene therapy. Normalized Δ NER score is represented by bar graph (bottom).

To translate the importance of macrophage activation to humans, we designed an inflammatory signature (Inflammatory score, Supplementary Table 3) and used it to interrogate two different data sets. First, we re-analysed a publicly available dataset of WAS iPSCs-derived macrophages (<u>GSE107963</u>, TPM normalized reads), and their isogenic counterparts that have been corrected to re-express WASp (Yuan et al., 2022). Despite high variations among samples, we found the expression of the inflammatory score is inversely correlated to WASp expression (Fig.6D). Remarkably, some genes within the NER signature (*Il6, Ccl8, Cxcl10 and Ctsb)* were consistently reduced by reintroducing WASp. The same trend was observed for other genes in the signature, albeit less consistently (Fig 6E). Moreover, those nuclear mechanics related genes controlled by WASp in mouse macrophages, were consistently lower in WASp mutated human macrophages (Fig.6F).

We also performed a bulk RNA-seq on PBMCs from a cohort of 6 WAS patients that were treated by gene therapy at SR-TIGET. Major WAS defects (i.e. BCR signalling and genes related to platelets function) were normalized post gene therapy (post-GT), without inducing alteration in other Arp2/3 activators (Supplementary Fig.6B and C). Interestingly, despite the complex composition of PBMCs that comprises several subsets (mainly B, T, NK and monocytes) and patients’ heterogeneity, we observed a consistent decrease in the inflammatory score post-GT (Fig.6G). Selected inflammatory genes emerging from RNA-seq, were validated by RT-qPCR, confirming that re-introducing WASp is sufficient to normalize inflammatory gene expression to the level of healthy donors (Supplementary Fig.6D). Intriguingly, the overall correction index was greater in samples showing a higher basal level of inflammation (Supplementary Fig.6E). In line with previous results, the NER signature score was elevated in five out of six patients, with some specific genes (Il6, *Ctsb, Ccl3, Cxcl10*) strongly depending on WASp expression (Fig 6H). These results support a role for WASp in restraining the inflammatory activation of human macrophages and, possibly, of other immune subsets.

## Discussion

Biophysical cues expressed in tissues influence the inflammatory functions of macrophages by a range of cellular mechanisms, including mechanotransduction at the nuclear envelope. In this study we show that WASp acts as a barrier to avoid nuclear rupture in mechanically challenged macrophages, restraining inflammatory responses. This direct connection between WASp-mediated actin polymerization and nuclear integrity has emerged from an experimental setup that recapitulates tissue forces experienced by cells *in-vivo.* Hence, this approach uncovered an essential supplementary function for the hematopoietic-specific actin nucleation-promoting factors WASp, beyond the well-established role at the cell cortex and in intracellular trafficking (Linder et al., 1999; Piperno et al., 2020; Gaertner et al., 2022). These findings are particularly relevant given the intrinsically migratory nature of immune cells and the nucleus as the main rate-limiting factor for constricted migration.

Previous studies showed that WASp null dendritic cells migrate faster inside microchannels or through non-deformable constrictions. However, the findings were not interpreted as dependent on altered nuclear properties (Gaertner et al., 2022; Oliveira et al., 2022). Our analysis confirms that WASp null macrophages, likewise dendritic cells, are facilitated in entering small constrictions. Additionally, we demonstrate that this is linked to an enhanced ability of the nucleus to deform, which correlates to a predisposition to undergo nuclear rupture. Cells harbouring defects in Lamin A/C or a disrupted nuclear-cytoskeleton coupling show similar nuclear deformations, increased migration and high rates of nuclear ruptures when exposed to mechanical confinement (Davidson et al., 2014; Harada et al., 2014; Kim et al., 2017; Bell et al., 2022). In line with our findings, a parallel study by our collaborators observed increased nuclear deformation and ruptures in confined dendritic cells that lack the WASp effector Arp2/3 (Delgado et al., 2024). More than one mechanism, not mutually exclusive, may explain reduced nuclear integrity in WASp null cells. The dense F-actin patches formed upon confinement and the immediate transcriptional response to supply factors controlling actin branching in WT, but not in WKO cells, suggest that WASp may induce cytoskeletal remodelling downstream of mechanical stimuli, creating a protective perinuclear actin barrier. Alternatively, as suggested by the presence of thin actin rings around the nucleus of a fraction of WKO cells and by gene expression data, lack of WASp may drive compensatory actin polymerization by different factors. As documented in a previous report, the resulting abnormal actin network may cause rupture by exerting excessive pressure on the NE under confinement (Mu et al., 2020). Interestingly, similar perinuclear actin rings were also observed in epithelial cells upon mechanical stretching, which has a role in promoting chromatin compaction (Le et al., 2016). Moreover, we cannot exclude that WASp limits nuclear rupture by acting inside the nucleus(Caridi et al., 2019), protecting it from DNA damage (Kovacs et al., 2023) or by mediating DNA repair dynamics via Arp2/3 (Delgado et al., 2024).

Of note, despite mild Lamin A/C defects and slight accumulation of DNA damage in WASp null macrophages, we found that the number of tissue-resident cells *in-vivo* was not affected, as opposed to the phenotype in Arpc4 KO and Lamin A/C deficient animals where Langerhans cells and alveolar macrophages, respectively, are depleted in tissues (De Silva et al., 2023; Delgado et al., 2024). Therefore, WASp ablation induces a proinflammatory state that does not culminate in cell death, suggesting a milder phenotype than complete blockade of Arp2/3 or Lamin A/C deletion. Whether this results from compensation by other NPFs remains to be investigated.

The unscheduled release of genomic DNA into the cytosol has been documented consequently to DNA damage (Dunphy et al., 2018; Mukherjee et al., 2019; Song et al., 2021), defective Lamin A/C (De Silva et al., 2023; En et al., 2024) and disrupted mechano-signalling due to reduced expression of Arp2/3, a direct WASp effector (Sladitschek-Martens et al., 2022). The latter study associated nuclear instability and nuclear envelope rupture to cGAS-STING activation and senescence in fibroblast. We observed a similar inflammatory activation downstream of NE rupture in WKO, raising many interesting points. While STING predominantly activates IRF3 upon exogenous introduction of DNA in the cytosol by transfection, leakage of genomic DNA following DNA damage leads to unconventional STING signalling, predominantly engaging NF-kB rather than IRF3 (Dunphy et al., 2018). Our NER inflammatory activation skewed towards cytokines rather than interferon supports the latter mechanism of STING engagement. Importantly, our work does not exclude the potential involvement of other DNA sensors nor the role of additional mechanisms connecting nuclear deformation and rupture to proinflammatory signalling in WASp null cells. Remarkably, the NER inflammatory signature induced in confined WASp null macrophages has been linked to the induction of senescence in fibroblasts (Sladitschek-Martens et al., 2022). Even if we have not explored senescence in WASp null cells or tissues, the expression of this signature under confinement, together with Lamin A/C defects and nuclear rupture, lead us to speculate that lack of WASp may predispose to accelerated ageing of the myeloid compartment *in-vivo*. Moreover, these findings suggest that inflammation may be connected to nuclear alterations in a broader group of diseases referred to as actinopathies (Papa et al., 2021; Sprenkeler et al., 2021), underscoring the relevance of investigating mechanical control of inflammation in immune cells. From a clinical perspective, our study has implications for understanding the molecular basis of WAS autoimmunity. Transcriptional data in human iPSCs-derived macrophages and total cells from the peripheral blood of gene therapy-treated WAS patients consistently indicate that WASp restricts the accrual of inflammation. However, inflammatory relapses are still frequent after treatment in WAS, including following hematopoietic stem cell transplantation (Sudhakar et al., 2021). According to recent evidence, including the data presented here, gene therapy may offer better control of inflammation in treated patients (Sereni et al., 2019; Magnani et al., 2022). Nevertheless, providing a further element to understand the origin of inflammation in WAS may help the management of residual disease, especially for those patients who do not have access to gene therapy.

In summary, our results unveil a cell-intrinsic WASp-mediated mechanism to control nuclear stability and inflammation under physiologically relevant settings, expanding the role of WASp-driven Arp2/3 actin polymerization in preserving myeloid cell homeostasis in tissues.

## Materials and methods

### Reagents and tools table

| Reagent/Resource | Reference or Source | Identifier |
| --- | --- | --- |
| **Experimental Models** |  |  |
| WASp<sup>-/-</sup> mice - B6.129S6-Wastm1Sbs/J | Jackson #:019458 | IMSR_JAX:019458 |
| HC and WAS PBMCs | San Raffaele Institute | Supplementary Table 5 |
| **Recombinant DNA** |  |  |
| pCDH-NLS-GFP | Addgene | #132772 |
| pMD2.G | Addgene | #12259 |
| psPAX2 | Addgene | #12260 |
| SEWW-eGFP-WASp | (Blundell et al., 2008) | / |
| **Antibodies Flow cytometry** |  |  |
| CD45 - APC-Cy7 | BioLegend | 103114 |
| CD11c - Bv786 | BioLegend | 117335 |
| F4/80 - Bv421 | eBioscience | 25-5870-80 |
| MHC-II - AF700R | BioLegend | 151507 |
| LY6C - PE | eBioscience | 11-0541-81 |
| CD64 - BV605 | BioLegend | 108923 |
| SiglecF - PerCP-Cy5.5 | BioLegend | 107105 |
| CD80 - APC | BioLegend | 139323 |
| CD86 - Bv650 | BioLegend | 104713 |
| CD3 - FITC | BioLegend | 100305 |
| LY6G - FITC | Bio-Rad | MCA6077GA |
| B220 - FITC | BioLegend | 103208 |
| CD19 - FITC | BioLegend | 115503 |
| NK1.1 - FITC | BioLegend | 116410 |
| **Antibodies FACS** |  |  |
| CD11c - Bv786 | BioLegend | 117308 |
| SiglecF - PerCP-Cy5.5 | BioLegend | 121406 |
| **Antibodies immunofluorescence** |  |  |
| Anti-Emerin | Thermo Fischer | PA5-29731 |
| Anti-H3K9me2 | Abcam | ab1220 |
| Anti-LMNA/C | Cell Signalling | mAb#4777 (4C11) |
| Anti-yH2AX | Sigma Aldrich | 05-636 (JBW301) |
| Anti-Mouse-488, Donkey | Invitrogen | A21202 |
| Anti-Mouse-555, Goat | Invitrogen | A21424 |
| Anti-Mouse-647, Chicken | Invitrogen | A21463 |
| Anti-Rabbit-488, Goat | Invitrogen | A21441 |
| Anti-Rabbit-594, Donkey | Invitrogen | A21207 |
| **Antibodies western blot** |  |  |
| Anti-GAPDH-HRP | Sigma Aldrich | G9295 (GAPDH-71.1) |
| Anti-WASp | Santa Cruz Biotech | sc-13139 (B-9) |
| Anti-Lamin A/C | Cell Signalling | mAb#4777 (4C11) |
| Anti-STING | Cell Signalling | 13647 (D2P2F) |
| Anti-TUBα-HRP | Cell Signalling | #9099 (11H10) |
| Anti-mouse-HRP | Jackson | 115-035-003 |
| Anti-Rabbit-HRP | Jackson | 711-035-152 |
| **Chemicals, Enzymes and other reagents** |  |  |
| 2-mercaptoethanol | Gibco | 31350010 |
| ACK lysis buffer | Gibco | A1049201 |
| Alexa Fluor™ 555 Phalloidin | Thermo Fischer | A34055 |
| Alt-R® Cas9 Electroporation Enhancer | IDT | 1075916 |
| Alt-R® CRISPR-Cas9 tracrRNA | IDT | 1072533 |
| Alt-R® Negative Control crRNA #2 | IDT | 1072545 |
| Alt-R™ crRNA STING | IDT | Mm.Cas9.TMEM173.1.AA |
| Alt-R™ crRNA WASp | IDT | Mm.Cas9.WAS.1.AC |
| Alt-R™ spCas9 Nuclease V3 | IDT | 1081058 |
| Bovine Serum Albumine (BSA) | Sigma Aldrich | A8806 |
| Clarity ECL substrate | Bio-Rad | 170-5060 |
| Collagenase type 2 | Worthington | LS004176 |
| Counting Bright beads | Miltenyi | C36950 |
| DMXAA | Invivogen | tlrl-dmx |
| DNase I from bovine pancreas | Millipore-Sigma | 11284932001 |
| EDTA | Thermo Fischer | 15575020 |
| FBS - low endotoxin | Euroclone | ECS0186D - lot. EUS037123 |
| Fibronectin Bovine protein | Thermo Fischer | 33010-018 |
| Gentamicin | Gibco | 15710064 |
| Glycine | Sigma Aldrich | G7126 |
| Hepes | Invitrogen | 15630080 |
| Hoechst 33342 | Invitrogen | H3570 |
| IMDM medium | Thermo Fischer | 31980030 |
| Lenti-X concentrator | Takara | 631231 |
| Live and Dead staining --Bv510 | Invitrogen | L34966 |
| LPS | Invivogen | tlrl-pb5lps |
| Mouse CD11c microbeads | Miltenyi | 130-125-835 |
| Mowiol 40-88 | Sigma Aldrich | 324590 |
| PDMS; SYLGARD 184 | Dow | 1673921 |
| PFA | Thermo Fischer | 28908 |
| Poly-Ethylene-Glycol (PEG) | Sigma Aldrich | 81260-1KG |
| PrimeScript™ RT Reagent Kit | Takara | RR036A |
| Protein inhibitor cocktail | Sigma Aldrich | P8340 |
| Recombinant mouse CCL19 (MIP3-β) | Gibco - Peprotech | 250-27B-1MG |
| Recombinant mouse MSC-F | Miltenyi | 130-101-706 |
| RIPA buffer | / | / |
| RNA Later™ | Invitrogen | AM7021 |
| RPMI 1640 medium | Thermo Fischer | 72400047 |
| Sodium Azide | Sigma Aldrich | 13412 |
| Sodium pyruvate | Gibco | 11360070 |
| Supernatant-containing GM-CSF | Produced by J558 cells | / |
| TB Green® Premix kit | Takara | RR420A |
| Triton-X-100 | Sigma Aldrich | X100-500ML |
| TRIzol™ reagent | Thermo Fischer | 15596026 |
| Tween-20 | Sigma Aldrich | 02286 |
| **Kits** |  |  |
| BCA assay kit | Thermo Fischer | 23225 |
| RNA high-sensitivity kit | Agilent | 5067-5579 |
| RNeasy micro kit | Qiagen | 74004 |
| Universal Index Twist kit 96 | Diatech-Labline | 101307 |
| WatchMaker - mRNA Library Prep Kit | Diatech-Labline | 7BK0001-024 |
| **Software** |  |  |
| Fiji (ImageJ) | v.1.54f | / |
| FlowJo | Tree Star, Inc. v.10.8.2 | / |
| Graphpad | Prism v.8.0.1 | / |
| R software | R version 4.3.1 | / |
| Biorender | www.biorender.com |  |

| Other |  |  |
| --- | --- | --- |
| J-558 cell line (GM-CSF producers) | / | / |

## Methods and protocols

### Mice

WASp^−/−^ mice (IMSR_JAX:019458) on a C57BL/6 genetic background were a gift from S. Snapper (Massachusetts General Hospital, Boston, Massachusetts, USA) (Snapper et al., 1998). Animal care and experimentation were conducted under national and international laws and policies (Italian D.L.gs. no. 26/2014 art.3.1A; European Directive 63/2010/UE). Mice were bred and maintained at the ICGEB animal Bio-experimentation facility in sterile isolators (12h/12h light/dark cycle, T 21 °C ± 2 °C, RH 55% ± 10%) and received a standard chow diet and water *ad libitum*. Experiments involving *ex-vivo* bone marrow culture were performed using WKO males’ mice and WT littermates as controls of 8-12 weeks of age. In case of *in-vivo* experiments WASp^-/-^ and heterozygous WASp^+/-^ female mice of 10 and 25 weeks of age were employed.

### Primary murine cell culture and human PBMCs

Mouse BM-derived macrophages (BMDMs) were generated in vitro from BM of C57BL/6 WT or WKO mice using recombinant M-CSF (rMCSF, Miltenyi Biotech). BM-derived cells were cultured at a concentration of 1 × 10^6^ cells/mL in non-treated 10 cm Petri dishes (Greiner) for 7 days using RPMI 1640 supplemented with 20 ng/mL rMCSF, 10% low endotoxin FBS (Euroclone), plus 50 μg/mL gentamicin (Gibco), 50 μM 2-mercaptoethanol (Gibco) and 1% Sodium Pyruvate (Gibco). Mouse BM-derived DCs (BMDCs) were generated in vitro from bone marrow of C57BL/6 WT or WKO mice using supernatant containing GM-CSF produced from J558 cell line. BMDCs were cultured at a concentration of 1.5 × 10^6^ cells/mL in Nunc-nontreated 6 well-plates (Thermo Fisher Scientific) for 7 days using IMDM supplemented with 30% GM-CSF supernatant, 10% low endotoxin FBS (Euroclone), 50 μg/mL gentamicin (Gibco) and 50 μM 2-mercaptoethanol (Gibco). All mice used for BM culture were males of 7 to 12 weeks of age.

Human PBMCs were gently provided by Dr. Anna Villa from San Raffaele Institute and shipped in RNA Later™ (Thermo Fischer Scientific). RNA extraction and library preparation for sequencing are described in the respective paragraphs. Information about patients is provided in Supplementary Table 4.

### Flow cytometry and sorting of alveolar macrophages

Alveolar macrophages (AMO) were obtained from homozygous WASp^-/-^ or heterozygous WASp^+/-^ female mice of 10 or 25 weeks old. After animal sacrifice, lungs were perfused in PBS to wash away red blood cells and digested with collagenase type 2 (265 U/mL; Worthington) and DNase I (250 U/mL; Millipore-Sigma) in complete media at 37°C for 30’. Collagenase activity was then stopped by EDTA 10 mM, and cell suspension was filtered using 100 μm cell strainer (Corning). The obtained single cell suspension was enriched for CD11c+ population using the mouse CD11c microbeads (Miltenyi) and stained with anti-CD11c, anti-CD45 and anti-SiglecF. 5 × 10^6^ cells/mL were resuspended in PBS 2% FBS for sorting and at least 100.000 AMO were sorted with a BD FACSAria II in PBS 20% FBS. Sorted AMO were centrifuged and frozen at -70°C in 350 μL of RLT lysis buffer prior to RNA extraction. Flow cytometry analysis was performed in a BD FACSCelesta™ starting from 1/5 of single cell suspension before CD11c enrichment, using antibodies reported in Supplementary Table 7 and Counting Bright beads (Miltenyi). All antibodies were incubated 1 h at 4°C in dark conditions. After antibody staining, samples were incubated 20’ at 4°C in dark conditions with Live and Dead fixable aqua (Invitrogen) 1:1000 in PBS and fixed over-night in PBS 1% PFA before a final wash and acquisition. Data analysis was performed on FlowJo software.

### RNA extraction and gene expression analysis

RNA from BMDM at steady state was obtained using of TRIzol^TM^ reagent (Thermo Fischer Scientific) and treated with DNaseI before retro transcription. RNA extraction and purification of sorted AMO, confined BMDM and human PBMCs was performed by RNeasy micro kit (Qiagen). Retro transcription and qPCR were done by PrimeScript™ RT Reagent Kit (Takara) and TB Green® Premix (Takara). Real time machine used for qPCR was a CFX96 Touch (Bio-Rad). Specific primers used for amplification were the following (5’-> 3’):

#### Mouse

Gapdh fwd AGAAGGTGGTGAAGCAGGCAT, Gapdh rev CGAAGGTGGAAGAGTGGGAGT; Il6 fwd GAGGATACCACTCCCAACAGA, Il6 rev AAGTGCATCATCGTTGTTCAT.

#### Human

Hprt1 fwd AATTATGGACAGGACTGAACGTCTTGCT, rev TCCAGCAGGTCAGCAAAGAATTTATAGC; Mxa fwd ACAGAACCGCCAAGTCCAAA, rev GCGGATCAGCTTCTCACCTT; Rsad2 fwd ATCCTTTGTGCTGCCCCTTG, rev TTGATCTTCTCCATACCAGCTTCC; Il6 fwd AGACAGCCACTCACCTCTTCAG, rev TTCTGCCAGTGCCTCTTTGCTG; Il1b fwd ATGATGGCTTATTACAGTGGCAA, rev GTCGGAGATTCGTAGCTGGA.

### Lentiviral preparation and BMDM infection

Second generation lentiviral particles were prepared by co-transfecting HEK-293T cells with plasmid of interest, relative packaging and envelope. 48 hours after transfection, supernatant-containing virus was collected, centrifuged 300g for 5’ to remove cell debris and filtered through 0.45 µm filter. Cleared supernatants were mixed with Lenti-X concentrator (Takara) and concentrated 100x, following manufacturer instructions. 20 µL of concentrated lentiviral particles were used to infect 2 x 10^6^ BMDM at day 4 in a 6-well plate. At day 6, viral-containing supernatant was removed and attached cells were washed 2x in sterile PBS. Infected BMDM were used at day 7 after two more washes in PBS to get rid of any remaining viral particle. Plasmid used for generation of lentivectors were the following: pMD2.G (Addgene #12259), psPAX2 (Addgene #12260), SEWW-eGFP-WASp (Blundell et al., 2008), pCDH-NLS-GFP (Addgene #132772), pTRIP-mCherry-FLAG-cGAS E225A/D227A (Addgene #127657).

### CRISPR-Cas9 RNPs electroporation

CRISPR-Cas9-mediated gene knock-out was achieved through electroporation of ribonucleoprotein (RNP) particles containing crRNA, tracRNA and Cas9 protein complexes. spCas9 and RNA oligos were purchased from IDT, annealed and mixed following manufacturer instructions. At day 4 of differentiation, 1.5 x 10^6^ cells were electroporated in 100 µL of media in a NEPA electroporator with the following parameters: poring pulse 275 V, 4 x 1 ms, interval 50 ms, D. Rate 10 %; transfer pulse 20 V, 5 x 50 ms, interval 50 ms, D. Rate 40 %, polarity +/-. After 24 h media was changed and cells were used for experiments at day 7 of differentiation.

### 6-well plate confiner

Mechanical confinement was performed in 6-well plates (Liu et al., 2015). 12 mm glass coverslips were washed with hand soap, rinsed in ddH_2_O, then in isopropanol and dried with a pressurized air gun. The clean coverslips were treated in a plasma cleaner (Diener Zepto) at 40W for 30 s. To make pillars of desired height, polydimethylsiloxane (PDMS; SYLGARD 184, Dow) was mixed at a ratio of PDMS A:cross-linker B 1:10 (wt/wt) and poured on top of wafer molds-containing pillars of 3 µm or 6 µm. The wafer molds, containing pillars of 3 µm or 6 µm, were fabricated by conventional micromachining from square pieces (25x25 mm) of a 500 μm thick silicon wafer. After cleaning the wafer with acetone and isopropanol, a positive photoresist MEGAPOSIT™ SPR™220 3.0 or 7.0 was spinned on. After the soft bake, they were exposed to a UV lamp for 30 seconds with a suitable mask and subjected a post-bake process. The development was carried out with MF24A. To functionalized with micropillars, glass coverslips were pressed to the wafers and baked at 85°C for 20 min before being carefully detached with a blade. After a wash in isopropanol, coverslips with micropillars were dried and plasma treated for 30 more seconds. Finally, these coverslips were washed in PBS and incubated with non-adhesive PEG solution (Sigma) 500 µg/mL in PBS for 30 min at RT and UV-sterilized for 20’. Before confinement, functionalized glass coverslips were washed in H_2_O and PBS and kept in sterile PBS at 4°C for a maximum of 48 h. To perform confinement experiments, we used large PDMS pistons placed on the plate lid to press the coverslips on the bottom of the plate. A drop of 250 µL containing 300.000 cells for immunofluorescent assays or 500.000 cells for RNA collection was confined for each well. 6-well plates with a glass bottom were used for cell imaging (MatTek, P06G-1.5-20-F); whereas non-treated 6-well plates (Corning) were adopted for RNA experiments. Nuclear blebbing quantification was performed right after confinement on live cells based on Hoechst 33342 staining (1:10.000 in PBS). Both types of confinements were kept for 1 h before fixation or RNA extraction. Fixation for IF was performed directly on confined cells, by leaving 4% PFA for 45 min before removing the lid, quenching PFA with 0,1 M glycine in PBS and washing cells 3x with PBS before proceeding with IF staining.

### Microchannel preparation and live imaging

Microchannels MC004 were purchased from 4DCell. And prepared following manufactures instructions. Coating was performed using 10 µg/mL fibronectin for 1 h at RT. Before utilization, channels were incubated from 30 min to overnight in 2 mL of BMDM media. Timelapse imaging was carried out overnight on a Nikon Eclipse TiE inverted microscope equipped with a 20x/0.75 NA dry objective, a motorized stage and a humidified chamber at 37°C and 5% CO_2_. Live imaging of migrating macrophages in microchannels was recorded with a frame every 5 min, based on NLS-GFP (FITC channel) and DIC (transmitted light). After overnight migration, cells inside channels were fixed by adding 10 µL of 4% PFA to each accession port and waiting for 1 h at RT before washing in PBS. Channels were kept in PBS 0,02% Sodium Azide until further imaging. Hoechst 1:1000 in PBS was added for at least 1 h before imaging fixed microchannels to facilitate detection and segmentation of nuclear morphology and herniations Fixed samples were imaged on a Zeiss LSM880 confocal microscope with a 63x/1.4 NA oil objective for individual nuclei and a 20x/0.60 NA dry objective for large field acquisitions and quantification of cell entry.

### Immunofluorescence imaging

Immunofluorescence staining at steady state was done with 50.000 BMDM seeded on fibronectin-coated (10 µg/mL, Millipore Sigma) 12 mm microscope slides (Epredia, Thickness value #1). Cells were left 30’ at 37°C and 5% CO_2_ to adhere. Fixation was performed in 4% PFA for 7 min at RT, followed by quenching in 0,1 M glycine in PBS for 3 min. After 3x washes in PBS, cells were permeabilized in PBS 1% BSA, 0,1% Triton-X100, blocked in PBS 3% BSA, 10 mM Hepes and stained ON at 4°C in primary antibodies diluted in PBS 0,2% BSA, 0,1% Triton-X100. Secondary antibodies were incubated at RT for 2h, followed by eventual phalloidin staining (Thermo Fischer Scientific, used 1:500) and 10 min nuclear staining with Hoechst 33342 (Invitrogen) 1:1000 in PBS. Slides were mounted with Mowiol and imaged at Zeiss LSM880 confocal microscope with a 63x/1.4 NA oil objective, or a 20x/0.60 NA dry objective. A full list of antibodies, dyes and probes used is presented in supplementary table 7.

### AFM based indentation measurements

AFM-based indentation measurements were performed by NanoWizard III AFM (JPK Instruments, Bruker) integrated with an inverted optical microscope (Axiovert 200; Zeiss) for sample observation. Cells were analysed in petri-dish and in culture medium at room temperature. The status of cells was constantly monitored by optical microscope. Indentation of the whole cell body was performed using MLCT-SPH-5um probes (Bruker), which have a cylindrical tip shank ending in a spherical tip with a 5 µm radius and a nominal spring constant of 0.06 N/m. Once localized the cell by optical microscope, the probe was brought over the nucleus area and pressed down to indent the cell. The motion of the z-piezo and the force were recorded. On each cell three to five force–displacement (F-D) curves were acquired with a force load of 0.5-1nN and at a rate of at 5µm/s in closed-loop feedback mode. Cell elastic properties were assessed by determining the Young’s modulus (E), which can be evaluated by fitting the force (F) indentation (δ) curves with the Hertz model.

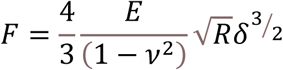

where F is the force, R is the radius of the spherical indenter and ν is the sample’s Poisson ratio (set to 0.5 for the cell) and δ is the indentation depth.

Data fitting was performed on the approaching F–D curves. JPK Data Processing software was used to convert the F-D curves into the force–indentation curves by subtracting the cantilever bending from the signal height to calculate indentation. The resulting force-indentation curves were fitted with the Hertz model for indentation depth of 500 nm.

### Transwell migration assay

24-well plates bearing transwell inserts of 5 µm and 3 µm. 5 µm plates were from Corning; 3 µm plates from Thermo Fischer Scientific. At day 7 of differentiation, cells were detached using MACS buffer (0,5% BSA, 2 mM EDTA in PBS), counted and resuspended in media without FBS containing 200 ng/mL of LPS (Invivogen) for 30 min in incubator. After LPS washout, cells were plated at a concentration of 200.000 cells in 200 µL for each insert in complete media. On the bottom of each well, 600 µL of complete media supplemented with 250 ng/mL of CCL19 (Peprotech) as chemoattractant were added to promote transmigration. Cells were kept 5 h in incubator to allow migration, following by collection of migrated cells for flow cytometry or IF analysis. Flow cytometry quantification of migrating cells was performed at BD Accuri™ C6 flow cytometer, using Counting Bright beads and normalizing the number of transmigrating cells to the total number of plated cells.

### Protein extraction and western blot

Cells were lysed in an appropriated volume of RIPA buffer (50 mM Tris/HCl pH 7.5, 150 mM NaCl, 1% NP-40, 0,1% SDS, 0,1% DOC in ddH_2_O) supplemented 1:30 with protease inhibitor cocktail (Sigma P8340). After centrifugation at 17000 g for 20 min, supernatant-containing protein extracts were kept at -20°C until further processing. Total protein concentration was quantified by means of BCA assay (Thermo Fisher), following manufacturer instructions. At least 20 µg of total protein lysates were loaded on a 10% custom made polyacrylamide gel and transferred on PVDF membranes upon SDS-page electrophoresis. Membranes were blocked at least 1 h at RT in 1x TBS, 0,1% Tween 20 (TBST), 3% dry fat milk. Primary antibodies were incubated ON at 4°C on a shaker. After 3x washes in TBST, secondary HRP-conjugated antibodies were kept 1-2 h at RT. Chemiluminescence signal was developed by Clarity ECL substrate (Bio-Rad) on a ChemiDoc™ Touch (Bio-Rad). A list of antibodies used is reported in supplementary table 7.

### Library preparation and RNA-sequencing

For mouse and human samples, total RNA was extracted using RNeasy Micro kit (Qiagen), according to the manufacturer’s protocol. RNA purity, integrity and quantity was checked at TapeStation 2200 (Agilent Technologies Inc., Santa Clara, CA, USA). RNA-seq libraries were prepared from 3 ng (human PBMCs) or 5 ng (mouse BMDM) of total RNA using mRNA Library Prep Kit (WATCHMAKER) following manufacture instruction. Briefly, we selected poly(A) using magnetic beads to focus sequencing on polyadenylated transcripts. After RNA fragmentation, a step of cDNA synthesis was performed. The resulting fragments were used for A-tailing and were ligated to the adapters and Index Anchors (TWIST bioscience, Twist universal system). PCR amplification (21 cycles) was performed to generate the final indexed cDNA libraries. Individual library quantification and quality assessment was performed using TapeStation 2200. Libraries were sent to Macrogen to be sequences on Novaseq (Human PBMCs) or NovaseqX (Mouse BMDM) instrument on Illumina platform as paired end run 150 bp reads. A total of 6 PBMCs samples were sequenced before as well after gene therapy. Four biological replicates of BMDM confined per genotype and three replicates for unconfined control were sequenced.

### RNA-sequencing analysis

Raw FastQ files were downloaded from Macrogen. Quality control, trimming and alignment were performed using standard pipelines for FastQC, Trimgalore and STAR.

#### Confiner data

Raw reads count matrix outputs were imported and analysed in R software (R version 4.3.1). Normalized reads were obtained by standard DEseq2 functions and used for heatmaps visualization. DESeq2 package (Love et al., 2014) was further used for differential gene expression analysis. GSEA was performed on gseGO function of ClusterProfiler package on Log2FC ordered genes with p-value cutoff of 0.01 and Benjamini-Hochberg p-value correction. DEGs were considered as genes bearing p-value< 0.05 and a Log2FC> 1.2 or Log2FC< -1.2. GO analysis was performed on DEGs using enrichGO function, based on biological processes, with a p-value cutoff of 0.05 and Benjamini-Hochberg statistical correction. Visualizations of functional enrichment results were obtained by ClusterProfiler and ggplot2 packages.

#### PBMCs data

DESeq2 analysis was performed starting on unnormalized reads comparing WAS pre- vs post-gene therapy samples. gseGO from ClusterProfiler have been used to perform GSEA on Log2FC ordered genes, using Entrez ID as key type, “BP” as ontology and 0.05 as p-value cutoff, without further statistical correction methods. Full list of GSEA is available as Supplementary Table 6.

#### Analysis of publicly available dataset from GSE107963

Data were downloaded as TPM normalized count matrix from GSE107963 (www.ncbi.nlm.nih.gov/geo/download/?acc=GSE107963, data deposited on 15.12.2022 and downloaded on 13.08.2024). Samples used for analysis refer to 3 WAS iPSCs (WASp -) and 3 isogenic iPSCs corrected to re-express WASp (WASp +), both differentiated into macrophages. TPM matrix was filtered and further processed to keep only genes with non-zero reads. Log2(TPM+1) was used for heatmap visualization.

### Image analysis

All raw images were analysed in Fiji ImageJ (Schindelin et al., 2012), using custom-written macros. <u>Lamin A/C alteration</u>: Lamin A/C alteration score has been calculated based on standard deviation (σ) of Lamin A/C signal. Nuclear segmentation has been based on Hoechst 33342 channel of z-projected nuclei, with a size threshold > 10 µm^2^, to exclude potential micronuclei from the analysis. Alteration score has been plotted as σ/10. Masks of Lamin A/C wrinkles for representative pictures has been obtained by an adaptive threshold method.

#### Chromocenters and H3K9me2 quantification

To quantify the number of chromocenters and the area of peripheral heterochromatin, nuclei were first segmented based on Hoechst 33342 signal. Chromocenters were detected based on an adaptive threshold and a size cutoff. Area of peripheral heterochromatin has been calculated as the sum of chromocenters’ area in close proximity to nuclear edges, as those within 1 µm from nuclear borders. The same cutoff has been used for H3K9me2 peripheral heterochromatin, measuring fluorescent intensity instead of chromocenters area.

#### Microchannels analysis and WASp-GFP distribution

Quantification of cells inside microchannels an nuclear rupture events have been conducted manually. Kymographs of NLS-GFP migrating cells were generated using a semi-automated macro in ImageJ. WASp-GFP distribution has been obtained in a semi-automated way as follow. Briefly individually cropped cells inside channels were subjected to automatic nuclear detection. Nuclear centre of mass was then moved to the centre of the image and all individual cells were cropped and horizontally aligned. The size of the cells was normalized at 600 pixels and expanded 50 times before saving at a resolution size of 1200x200 pixels (w x h). Size normalized cells were then merged by Stack>Z project>Average Intensity to obtain an average intensity map profile of WASp distribution.

### Statistics

Primary data were collected in Microsoft Excel and statistical analyses were performed using GraphPad Prism version 8. For comparison between two or more groups with normally distributed data two-tailed Student’s t-test, one-way ANOVA or two-way ANOVA were used as appropriate. The non-parametric Kruskal-Walli’s test with Dunn’s multiple comparison was performed to compare three or more unmatched groups. In case of normalized values to 1 or 0, statistics has been calculated by a one-sample t-test. *P* values <= 0.05 were considered significant and marked by asterisk, if not stated differently.

## Data availability

Sequencing data have been deposited at GEO database: BMDM confiner data (accession: GSE285024), publicly available as of the 24/12/2024 and PBMCs WAS data (accession: GSE285174), publicly available as of the 25/12/2024. Source data are provided in supplementary files. Any additional information required to reanalyse the data reported in this paper is available from the lead contact upon formal request.

## Author contributions

**R.A**, **G.M.P** and **F.B** conceived the study and designed the experiments. **R.A** performed most of the experiments, data analysis, and prepared the figures. **R.A** and **F.B** wrote the manuscript. **G.B** helped in the execution of experiments and performed WASp-GFP confinement. **Z.A** provided training and support for the confiner assays and helped in performing confiner experiments in Fig.3 (A, B and C). **M.M** assisted in image analysis. **G.M.P** and **L.L.R** assisted in the experimental part and critical discussion of the data. **G.M.P** prepared library for sequencing. **M.C** prepared photolithographic molds used for confiner slides fabrication. **L.A** performed and analysed atomic force microscopy. **M.C.C**, **F.F**, **A.A** and **A.V** provided human PBMCs samples. **A.-M.L.-D** critically discussed the data and the manuscript. **F.B** supervised the whole study. All authors read and approved the final version of the manuscript.

## Acknowledgements

This work was supported by Telethon grant GGP20102 to **F.B**. **G.B**, **G.M.P** and **L.L.R**-were supported by ICGEB Arturo Falaschi pre- and post-doctoral fellowships. **R.A** was supported by Italian Telethon. **R.A** received an EMBO scientific exchange grant (10017) to perform part of the work in the lab of **A.-M.L.-D**. We Thank Luca Triboli from ICGEB Trieste to help discussing bioinformatic data analysis.

---

**Supplementary Figure 1.**
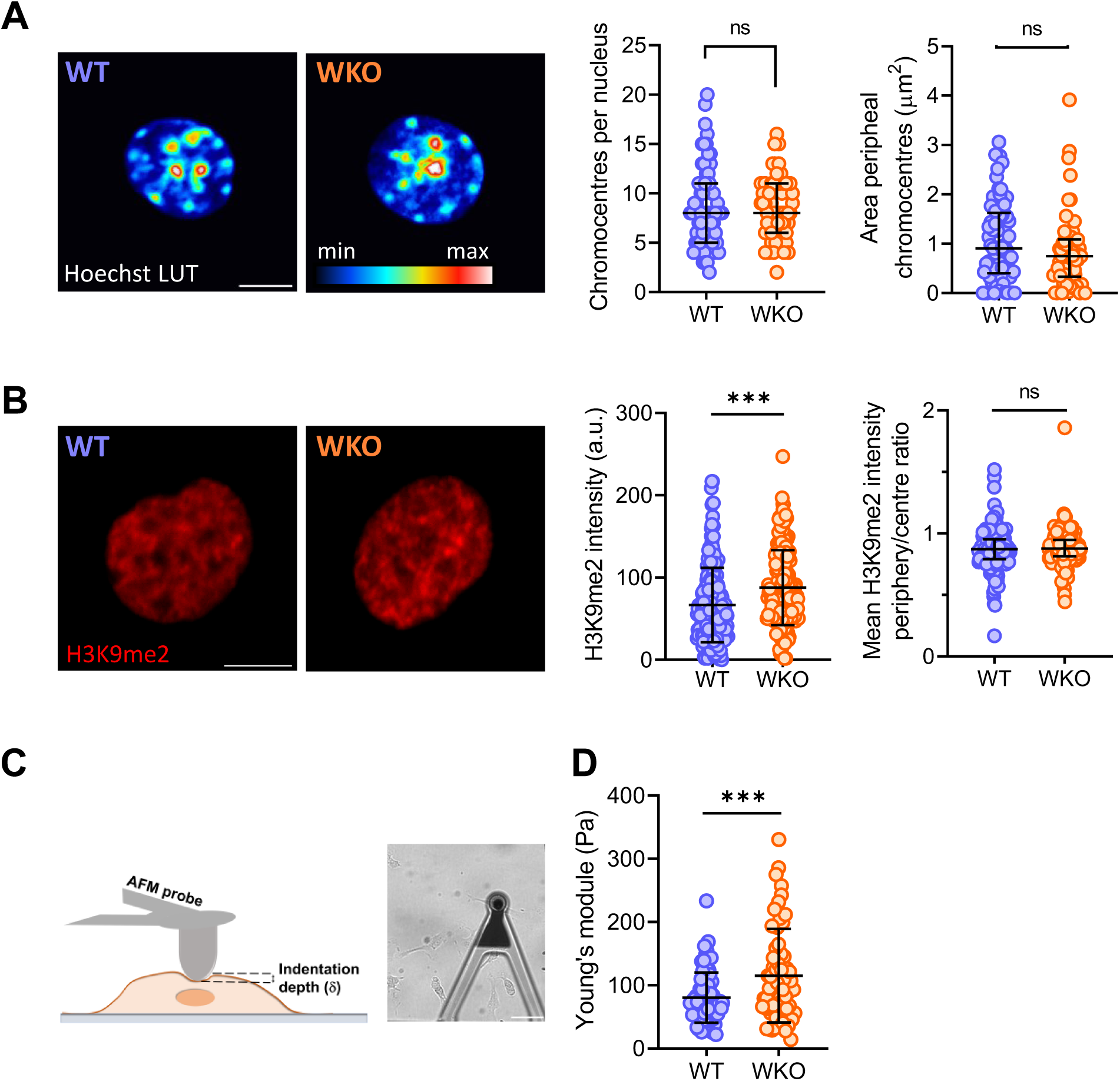
Chromatin features and cellular mechanics in WT and WASp KO macrophages. (**A**) Representative images (left) and quantification (right) of the number of chromocenters (heterochromatin foci) in WT and WKO BMDM nuclei and the mean area of peripheral heterochromatin foci. *n*=65 individual nuclei measured from *N*=3 independent experiments. Scale bar 5 µm. Statistics was calculated by two-tailed unpaired t-test. ns, not significant. (**B**) Representative images (left) and quantification (right) of mean fluorescent intensity and ratio of peripheral vs central H3K9me2 marker of facultative heterochromatin. *N*=3 independent experiments, with *n*=65 individual nuclei measured in total. Data were analysed by two-tailed unpaired t-test. \*\*\**P*<=0.001; ns, not significant. Scale bar 5 µm. (**C**) Scheme of AFM-indentation measurement of a single BMDM cell. Bright field image scale bar 50 µm (**D**) Quantification of Young’s modulus of *n*=71 WT and *n*=62 WKO BMDM cells, from three independent measurements. Data were analysed by two-tailed unpaired t-test. \*\*\**P*<=0.001.

**Supplementary Figure 2.**
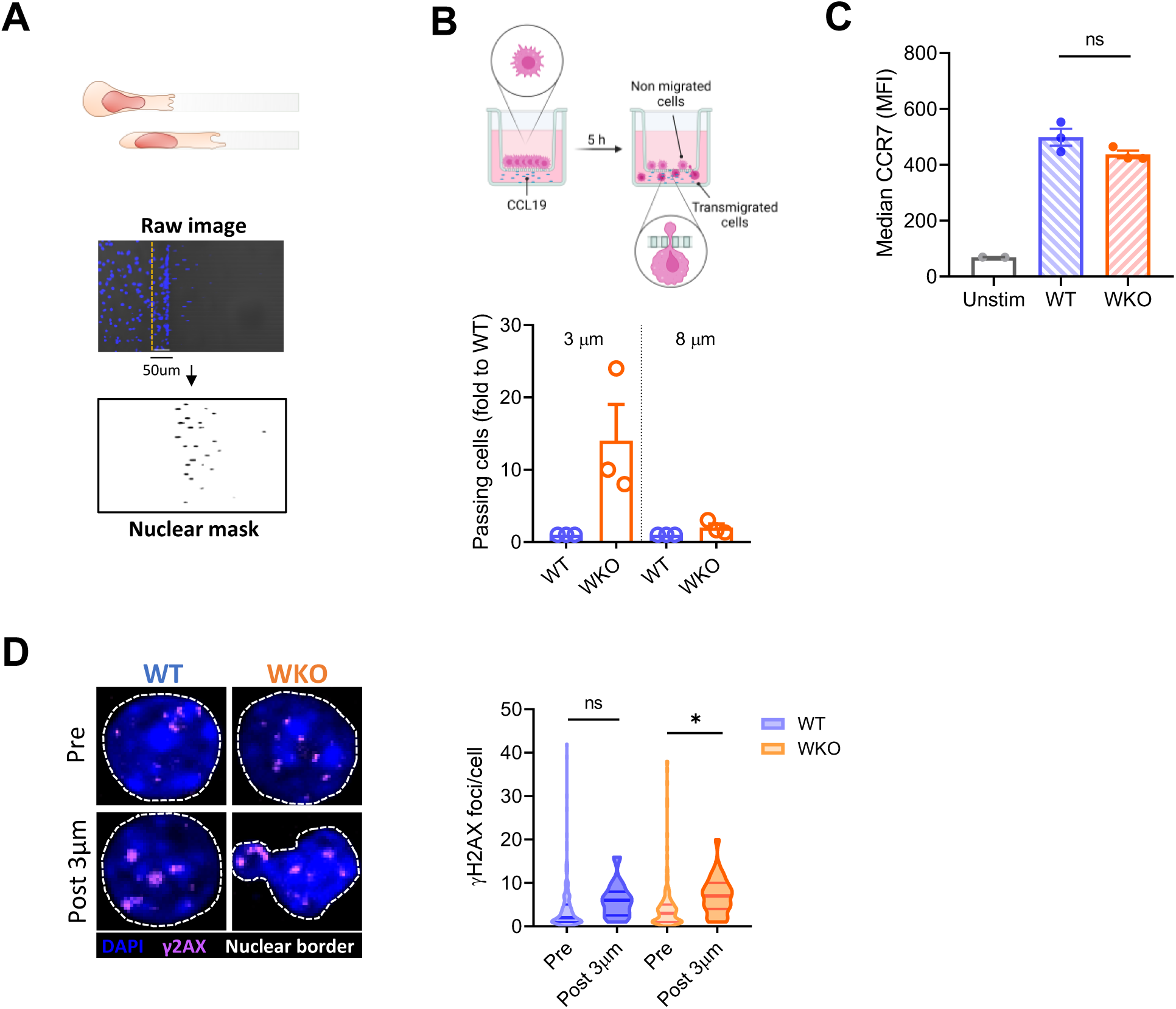
Constricted migration augments DNA damage in WKO cells. (**A**) Schematic overview of nuclear mask segmentation from raw channel images and quantification of entrance rate as percent of cells inside channels and approaching cells. Approaching cells were defined as those present within 50 µm from channel beginning (yellow dashed line). (**B**) Schematic overview of transwell assay used for BMDCs migration (top) and relative quantification of transmigrating cells (bottom), in transwell of indicated pore size. *N*=3 independent replicates are plotted as normalized fold to respective WT (value=1). Statistics was calculated using a one-sample t-test. (**C**) Representative pictures of yH2AX staining pre and post 3 µm transwell migration in BMDCs and relative quantification of number of yH2AX foci per cell in WT and WKO. Data are from *N*=3 independent experiments with *n*=178; 202 (WT and WKO pre transwell) and *n*=13; 33 (WT and WKO post transwell). \**P*<=0.05 in a standard one-way ANOVA test. (**D**) Flow cytometry quantification of CCR7 median expression in WT and WKO BMDCs before 3 µm transwell migration and following 200 ng/mL of LPS stimulation for 30’ (*N*=3) or in unstimulated control (*N*=2). Statistics was performed comparing WT and WKO after LPS stimulation by an unpaired t-test. ns, not significant.

**Supplementary Figure 3.**
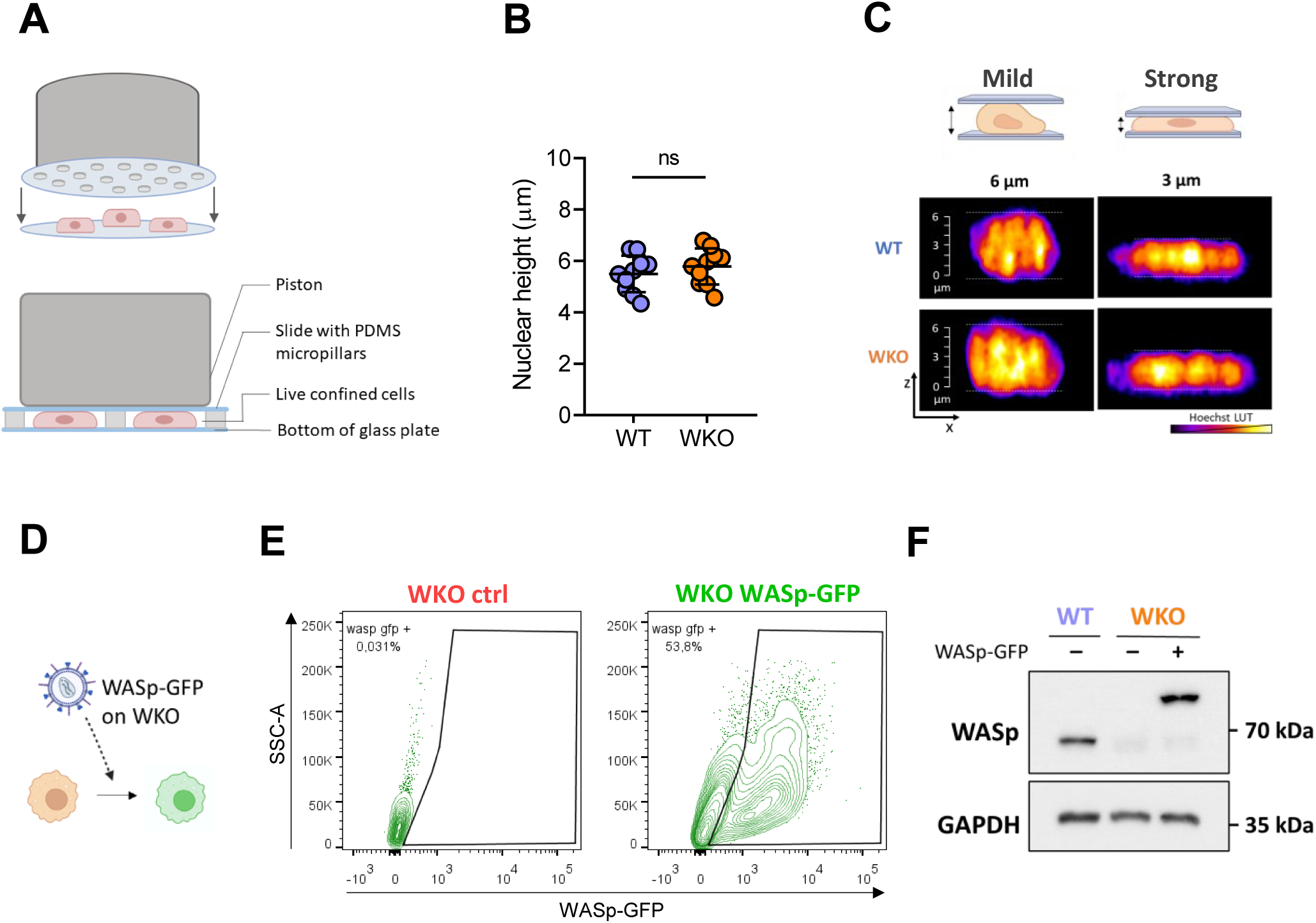
WASp limits nuclear rupture under confinement. (**A**) Cells were subjected to vertical confinement, as schematically represented. (**B**) Nuclear height quantification in 2D without confinement in BMDM. Data from *N*=1 experiment with a total of 10 z-stacks nuclei measured per genotype. (**C**) Representative images of BMDM nuclear height from live-imaging confocal acquisition of cell under confinement at 6 µm and 3 µm. Vertical scale bar 6 µm, with a tick every 1 µm. (**D**) WKO BMDM were transduced with lentiviral particles encoding WASp-GFP. (**E**) Flow cytometry shows the fraction of WKO cells expressing WASp-GFP to visualize WASp-GFP expression. (**F**) Representatives immunoblot of WASp expression in WT, WKO ctrl and WKO WASp-GFP macrophages after 7 days of differentiation.

**Supplementary Figure 4.**
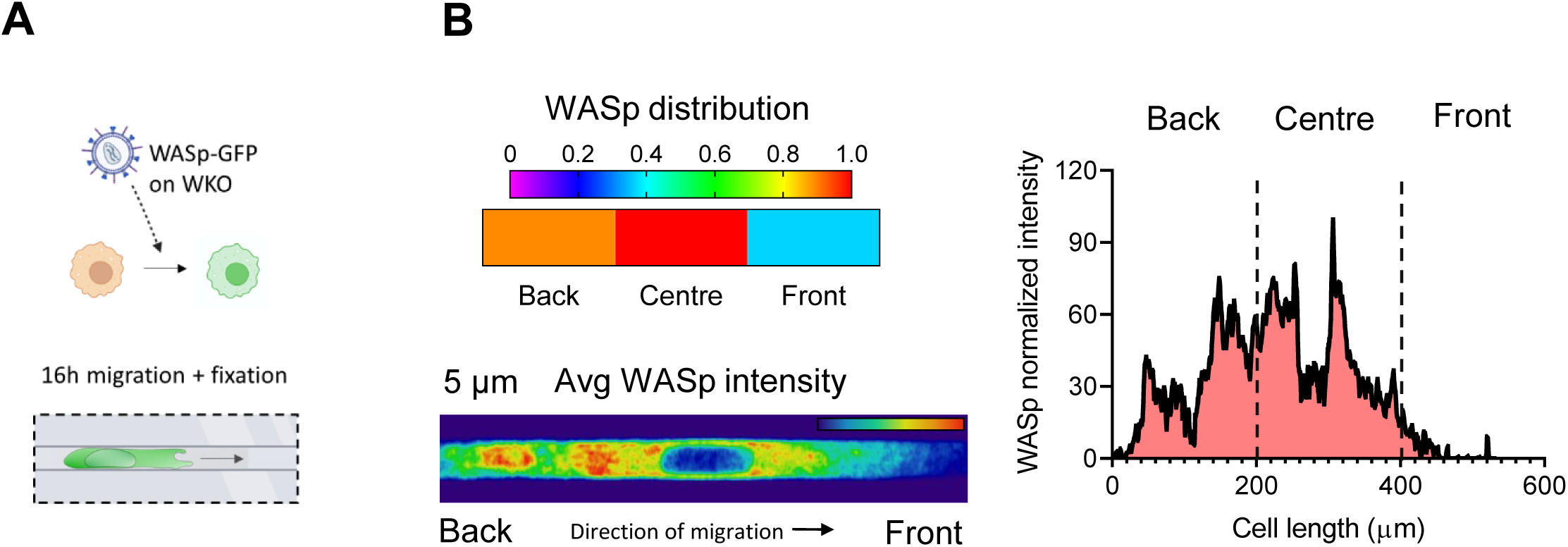
WASp perinuclear localization under confinement. (**A**) WASp-GFP expressing cells were recorded during migration in 5 x 5 μm microchannels. (**B**) After fixation at the end of the recording period, WASp distribution was calculated by dividing the cell volume into three equal parts (top left). Average intensity map of WASp-GFP was obtained by normalizing and merging different cells from a fixed end time point analysis (bottom left). Data are from *n*=42 cells obtained in *N*=2 independent experiments. Representative line intensity profile of WASp-GFP signal (right).

**Supplementary Figure 5.**
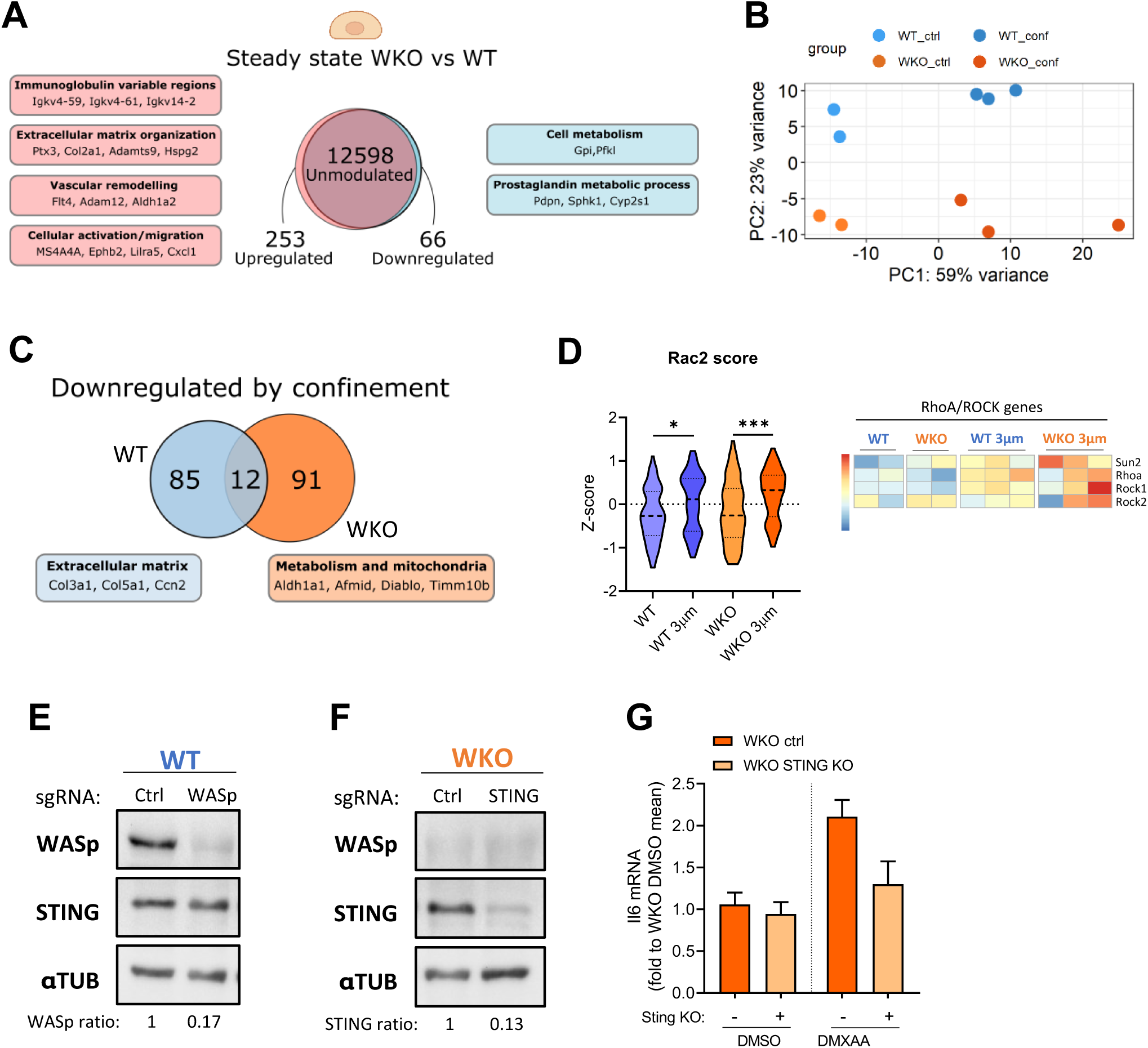
Mechanical confinement triggers innate activation in WASp KO macrophages. (**A**) Venn diagram of differentially expressed genes between WT and WKO in unconfined resting condition. Relevant genes and processes are highlighted. (**B**) PCA analysis of BMDM unconfined controls and 3 µm confined samples from RNA-seq. (**C**) Venn diagram of DEGs downregulated by confinement in WT and WKO at 3 µm. Relevant genes are highlighted. (**D**) Z-score of Rac2 signature across the two genotypes in unconfined and confined conditions (left). Normalized expression of Sun2/RhoA/ROCK genes is showed as heatmap (right). (**E**) Representative western blot of KO efficiency control in WT macrophages after 72 h from sgCtrl or sgWASp CRISPR-Cas9 electroporation. (**F**) Representative western blot for KO efficiency in WKO macrophages after sgCtrl or sgSTING CRISPR-Cas9. (**G**) Il6 quantification in WKO and WKO STING KO at steady state and upon direct STING stimulation by 3 µg/mL of DMXAA.

**Supplementary Figure 6.**
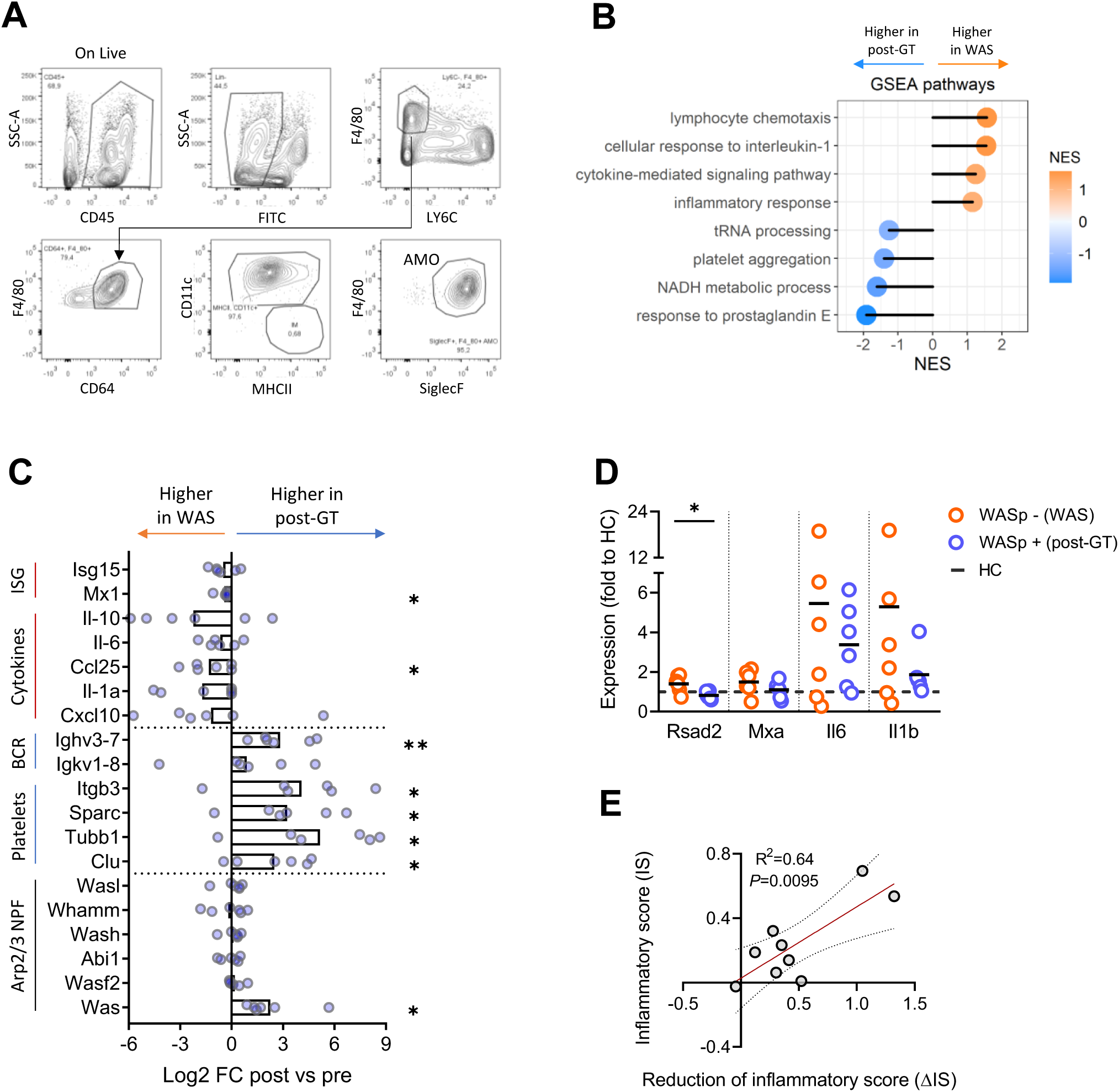
WASp reduces steady state inflammatory activation of macrophages in-vivo and in-vitro. (**A**) Flow cytometry gating strategy to identify AMO in mouse lung tissues. (**B**) GSEA analysis of relevant modulated pathways in WAS PBMCs before and after gene therapy correction. Values are expressed as normalized enrichment score. (**C**) Expression of the indicated genes in individual PBMC samples pre- and post-gene therapy. Data are plotted as Log2 FC post vs pre gene therapy, each dot is a patient. One sample t-test; \**P*<=0.05 and \*\**P*<=0.01. (**D**) RT-qPCR validation of selected inflammatory genes in 6 WAS patients pre- and post-gene therapy. The dotted line represents the mean gene expression of 3 healthy donor controls (HC). One sample t-test: \**P*<=0.05. (**E**) Correlation between inflammatory score (IS) before WASp restoration (y-axis) and IS reduction (Δ IS) upon WASp re-expression (x-axis). Data combines 3 human iPSCs-derived macrophages and 6 human PBMCs. Data were analysed by linear regression; *P*=0.0095.

